# Integrating Genomics, Collections, and Community Science to Reveal Speciation in a Variable Monitor Lizard (*Varanus tristis*)

**DOI:** 10.1101/2023.10.03.560725

**Authors:** Carlos J. Pavón-Vázquez, Alison J. Fitch, Paul Doughty, Stephen C. Donnellan, J. Scott Keogh

## Abstract

—The accurate characterization of species diversity is a vital prerequisite for ecological and evolutionary research, as well as conservation. Thus, it is necessary to generate robust hypotheses of species limits based on the inference of evolutionary processes. Integrative species delimitation, the inference of species limits based on multiple sources of evidence, can provide unique insight into species diversity and the processes behind it. However, the application of integrative approaches in non-model organisms is often limited by the amount of data that is available. Here, we show how data relevant for species delimitation can be bolstered by incorporating information from tissue collections, museum specimens, and observations made by the wider community. We show how to integrate these data under a hypothesis-driven, integrative framework by identifying the processes generating genetic and phenotypic variation in *Varanus tristis*, a widespread and variable complex of Australian monitor lizards. Using genomic, morphometric (linear and geometric), coloration, spatial, and environmental data we show that disparity in this complex is inconsistent with intraspecific variation and instead suggests that speciation has occurred. Based on our results, we identify the environmental factors that may have been responsible for the geographic sorting of variation. Our workflow provides a guideline for the integrative analysis of several types of data to identify the occurrence and causes of speciation. Furthermore, our study highlights how community science and machine learning—two tools used here—can be used to accelerate taxonomic research.

---

The mischaracterization of species diversity poses a major challenge to our understanding of the ecology, evolution, and conservation of biodiversity (Bini et al. 2006; Hortal et al. 2015). Therefore, generating accurate species delimitation hypotheses is a crucial goal of systematic biology. The rapid accumulation of large molecular datasets for non-model organisms has proven to be a valuable tool in this quest. On one hand, high-throughput sequencing may allow the detection of cryptic species in cases where morphology or smaller molecular datasets fail to provide evidence for speciation (Weiss et al. 2018). On the other hand, wider genome representation may reveal complex gene flow scenarios that would remain undetected by other types of data on their own, preventing the overestimation of species diversity (Hinojosa et al. 2019; Chan et al. 2020).

While large molecular datasets provide a plethora of information, there are caveats that need to be taken into consideration when using them for species delimitation. With increasing data there is the potential to detect finer-scale population structure, which can be misidentified as species-level divergence by some species delimitation approaches (Sukumaran and Knowles 2017). Isolation by distance (IBD)—the positive correlation between genetic and geographic distance—is common in natural populations and can confound species delimitation (Mason et al. 2020). Isolation by environment (IBE) may also misguide species delimitation and could be even more common than IBD (Sexton et al. 2014). Gene flow and incomplete lineage sorting are other common sources of uncertainty and conflict in species delimitation (Weber et al. 2019; Chan et al. 2020).

One way to minimize the impact of the shortcomings of the different methods and data is to implement integrative species delimitation, analyzing multiple sources of evidence in different ways. The implementation of integrative taxonomy often results in conservative taxonomic frameworks if species status is reliant on the congruence between data and approaches (Carstens et al. 2013). In this context, museum specimens remain a valuable source of information. Specimens convey information about the genotype, gene interactions, epigenetic regulation, and environmental demands in the form of morphological data. The approaches used to collect and analyze morphological data have advanced in parallel to molecular methods, providing unique insight into species limits (Pavón-Vázquez et al. 2018, 2022; Chaplin et al. 2020). Furthermore, museum specimens link specific phenotypes (and genotypes, if molecular resources are available) with particular temporal, geographic, and environmental conditions (Edwards et al. 2005; Schmitt et al. 2019). In that way, they can provide valuable insight into the drivers of genotypic and phenotypic divergence. Thus, the development of approaches that can integrate multiple sources of evidence to inform taxonomy (e.g., Solís-Lemus et al. 2015; Pyron 2023) is critical.

Compared to molecular data and museum specimens, observations made by the general public remain a largely underutilized resource in taxonomic research. Biodiversity community science projects and observations are accumulating rapidly (Pocock et al. 2017; Rowley et al. 2019; Messaglio and Callahan 2021), generating valuable data for understudied taxa and regions. Community observations—with iNaturalist (https://www.inaturalist.org/) as their best known repository—have been mainly leveraged in the context of conservation (McKinley et al. 2017; Chowdhury et al. 2023b). However, these observations also convey important information about the geographic and temporal distribution of morphological variation (Drury et al. 2019; Hantak et al. 2022). In that way, community science can provide important information about evolutionary processes and inform taxonomy (Fritz and Ihlow 2022; Fritz et al. 2023; Pizarro et al. 2023).

Morphologically variable species complexes represent a challenge for traditional species delimitation approaches. Geographic patterns of morphological variation may lead to overestimation of species diversity, or prove too complex to be easily translatable into binomial nomenclature. Molecular data can also result in inaccurate delimitations, especially if geographic clusters are generally concordant with phenotypic variation and the patterns of gene flow/isolation are not adequately characterized. The monitor lizard *Varanus tristis* (Fig. 1a) is a variable species complex with one of the widest distributions among Australian reptiles. Its range encompasses most of the continent, except for the coldest regions in the south (Figs. 1b, c) (Pianka 1971, 2004). The species exhibits considerable variation throughout its range, leading to the recognition of two subspecies: *V*. *tristis tristis* and *V*. *tristis orientalis*. These subspecies are mostly defined by coloration, with the former applied to dark-headed individuals and the latter to light-headed individuals (Pianka 2004; Cogger 2014). Dark-headed individuals are found in the central and western portions of the range.

**FIGURE 1.**
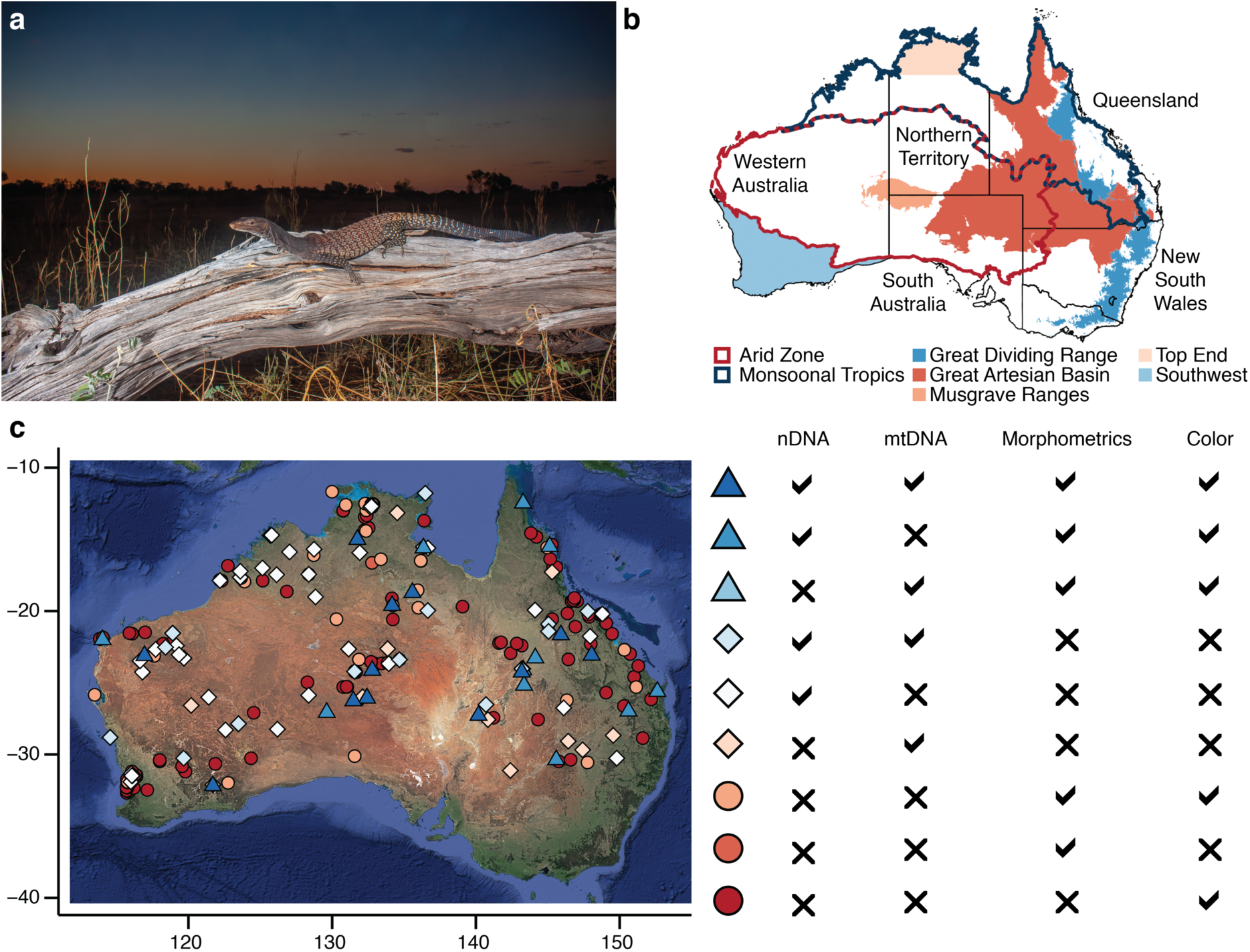
Focal taxon and study area. a) Adult male *Varanus tristis* from central Queensland, Australia; photograph by Kieran Palmer. b) Mainland Australia; black lines indicate state borders, with names next to them; other regions mentioned throughout the text are indicated by the colored polygons. c) Sampling of *V*. *tristis*; symbols indicate the major types of data available for each individual (circles = phenotypic only, diamonds = molecular only, triangles = both); colors indicate the specific types of data available; base map ©2023 Google.

Their bodies show varying degrees of melanism, with darker colors generally occurring in the southern portion of their range (Pianka 2004; Schmida 2020). Light-headed individuals are known from the northern Monsoonal Tropics and adjacent areas in eastern Australia (Pianka 2004; Schmida 2020). These populations show variation in the extent and conspicuousness of ocelli, as well as in head color (ranging from brownish-grey to yellowish/reddish). Individuals from the eastern portion of the range show a unique combination of coloration, scutellation, and morphometric traits (Storr 1980; Pianka 2004; Schmida 2020). This has led some authors to reserve the use of *V*. *t*. *orientalis* for this population (e.g., Storr 1980; Schmida 2020). In contrast, the Australian Society of Herpetologists Official List of Australian Species (2022) does not recognize subspecies within *V*. *tristis* because of a lack of reciprocal monophyly between light and dark-headed individuals in a mitochondrial study (Fitch et al. 2006) and the morphological intergradation between them (Pianka 2004). Ultimately, taxonomic clarity in these monitors is hindered by the lack of any comprehensive analysis of genetic and morphological variation.

Here, we present an integrative workflow to delimit species that capitalizes on the wealth of samples and information that are available in public repositories (Fig. 1c). Particularly, it uses a combination of molecular, environmental, and phenotypic data obtained from tissue samples, museum specimens, and community observations. The data are analyzed both iteratively (i.e., in separate steps) and integratively (i.e., simultaneously) to identify evolutionarily independent lineages. We demonstrate the utility of this workflow for species delimitation in taxonomically challenging complexes by applying it to *V*. *tristis*. Our analyses enabled us to distinguish between intraspecific variation and differentiation due to speciation.

## Materials and Methods

Our workflow has characteristics of both iterative and integrative species delimitation (Fig. 2). The procedure accounts for confounding factors such as IBD, IBE, incomplete lineage sorting, introgression, and local adaptation/plasticity. Furthermore, we specify our hypotheses and expectations under different scenarios at several points throughout the workflow. Additional details are provided in the following paragraphs.

**FIGURE 2.**
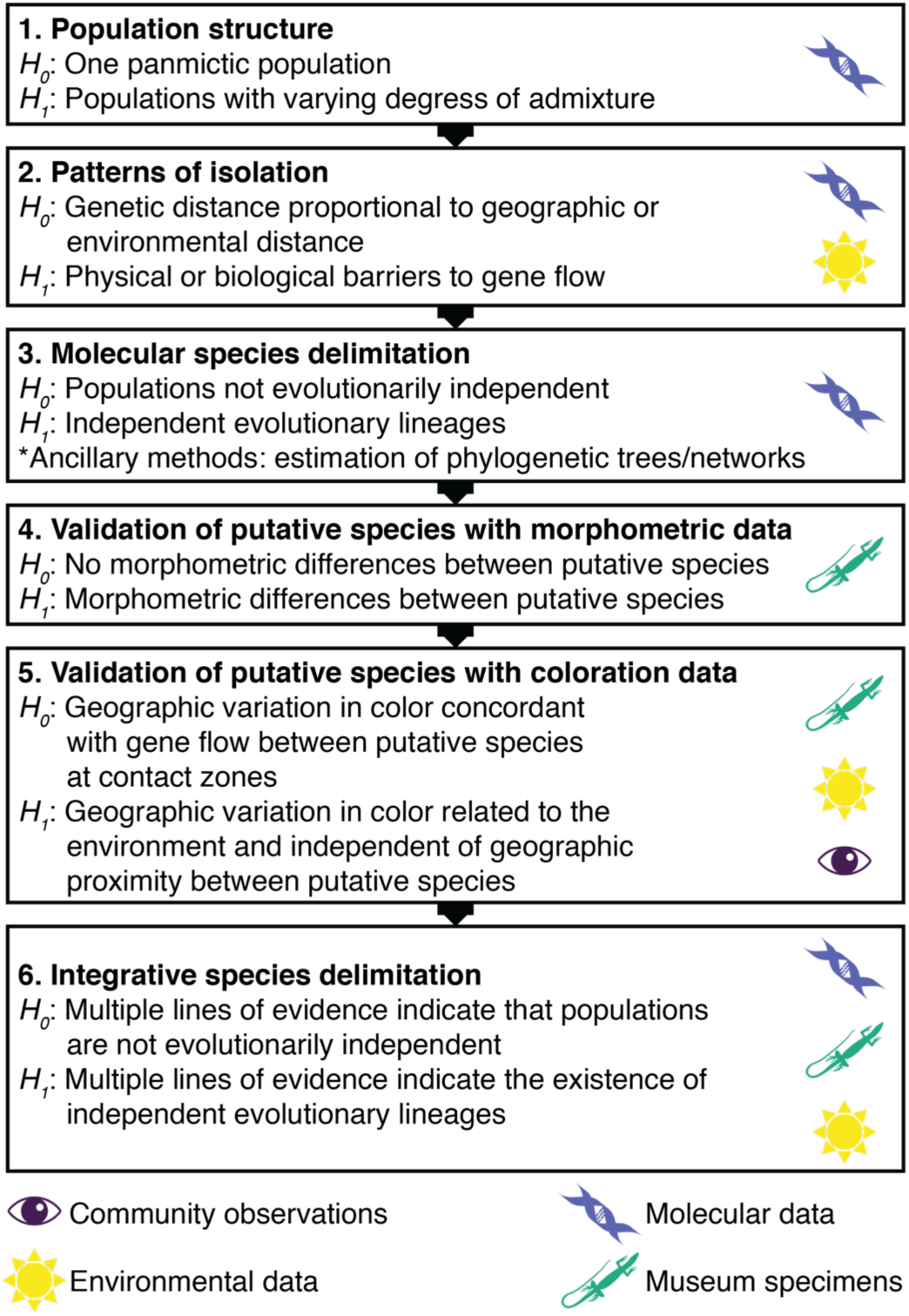
Integrative workflow implemented here to delimit species in the *Varanus tristis* species complex. For each step, the expectations under a scenario where no speciation has occurred (*H_0_*, in its broad sense) are indicated. Alternatively, evidence for speciation may be recovered (*H_1_*). In the third step, methods that are not specifically designed to distinguish between these scenarios but that can provide insight into the listed evolutionary processes are specified as ancillary methods. The sources of evidence used in each step are indicated by the icons.

### Molecular Sampling

We used the DArTseq (Diversity Arrays Technology [DArT], Canberra, Australia) platform to genotype 93 individuals from across the range of *V*. *tristis* (Fig. 1c). To root trees and test the monophyly of *V*. *tristis*, we also sequenced samples of *V*. *glauerti* (two individuals), *V*. *scalaris* (six individuals), and *V*. *varius* (two individuals). Tissue samples were sourced from several Australian biological collections (Table S1). DArTseq data have been used successfully for phylogeographic inference and species delimitation in numerous taxa (e.g., Georges et al. 2018; Chaplin et al. 2020; Esquerré et al. 2021; Binks and Byrne 2022; Mahony et al. 2022), including other *Varanus* (Pavón-Vázquez et al. 2022).

DArT’s proprietary pipelines were used to generate data for two primary sets of individuals: one including *V*. *tristis* only and one that also included the outgroups. DArTseq achieves complexity reduction through digestion with restriction enzymes and size filtering followed by sequencing (Sansaloni et al. 2010; Kilian et al. 2012). The enzymes used for this project were PstI and SphI. Paired-end sequencing was performed in an Illumina Hiseq2500 sequencer. Common prokaryotic contaminants were removed and the alignments were annotated based on the genome of *V*. *komodoensis* (BioProject PRJNA523222; Lind et al. 2019). The datasets delivered by DArT comprised 296,925 and 277,205 loci for the sets containing *V*. *tristis* only and the one that includes the outgroups, respectively. Sequences for each locus are short (∼69 bp) and contain at least one single-nucleotide polymorphism (SNP). We conducted additional filtering of loci and individuals on each dataset to ensure data quality and meet the assumptions of the different analyses (Table S2). Most processing was done in R 3.6.2 (R Core Team 2019) with the ‘dartR 1.8.3’ package (Gruber et al. 2018).

To detect potential mito-nuclear discordance, we sequenced the mitochondrial NADH dehydrogenase subunit 4 protein (*ND4*) gene and the associated *tRNA-His*. We generated data for 44 individuals following the protocols described by Fitch et al. (2006). The purified PCR products were sequenced on an ABI Prism® 3700 DNA Analyzer at The Institute of Medical and Veterinary Science (IMVS), Adelaide, South Australia. We complemented these data with 32 sequences obtained from GenBank (Table S1). We aligned the sequences using the MUSCLE algorithm (Edgar 2004) and refined them by hand in AliView 1.28 (Larsson 2014).

The final alignment comprised 663 bp and included the following species (number of individuals in parentheses): *V*. *tristis* (45), *V*. *glauerti* (3), *V*. *glebopalma* (2), *V*. *hamersleyensis* (7), *V*. *mitchelli* (2), *V*. *pilbarensis* (7), *V*. *scalaris* (7), *V*. *semiremex* (2), and *V*. *spenceri* (1).

### Population Structure

These analyses were performed on a filtered version of the *V*. *tristis* SNP dataset containing 81 individuals and 35,162 unlinked SNPs. First, we estimated the ancestry coefficients of each individual to detect deviations from panmixia and how they relate to geography. This allowed us to identify operational units for downstream analyses. Ancestry coefficients were calculated with sNMF (Frichot et al. 2014) in the ‘LEA 2.8.0’ (Frichot and François 2015) R package. sNMF is an efficient and accurate method based on sparse nonnegative matrix factorization (Frichot et al. 2014). We selected the combination of the regularization (*α*) and tolerance (*ε*) parameters that minimizes cross-entropy between 18 possible combinations. For each combination, we performed ten runs for each value of *K* (number of populations) between one and ten. Subsequently, we performed 100 runs each for *K* 1–10 using the selected combination of *α* (1) and *ε* (0.00001). We extracted the ancestry coefficients from the run yielding the lowest cross-entropy. Additionally, we used principal component analysis (PCA) to visualize the genetic similarity between individuals in two-dimensional space. This was done with the “gl.pcoa” function of ‘dartR’.

### Patterns of Isolation

We performed several analyses to test whether genetic distance between individuals increases gradually with geographic (IBD) and/or environmental distance (IBE). The detection of IBD and/or IBE would be indicative of a gradient of genetic differentiation within a single species. Alternatively, abrupt changes in genetic distance over geographic/environmental space are indicative of physical or biological barriers to gene flow.

We tested for IBD using a Mantel test as implemented by the “gl.ibd” function of ‘dartR’. We used the *â* statistic (Rousset 2000) as our measurement of genetic distance, since it may be more appropriate than *F_ST_*-based statistics when data points represent individuals instead of populations. Geographic distances were based on the Mercator projection. To locate geographic regions where genetic similarity decays abruptly, we estimated the effective migration surface of *V*. *tristis* in EEMS 0.0.0.900 (Petkova et al. 2016). This is a model-based Bayesian approach that identifies deviation from IBD. We used average genetic dissimilarity as our measurement of genetic differentiation (Petkova et al. 2016) and divided mainland Australia into 500 demes. We conducted three independent runs consisting of 1.5 million burn-in iterations, 5 million post-burn-in iterations, and a thinning of 10,000. The runs were combined after assessing for convergence.

To quantify the relative contributions of geographic and environmental distance to genetic differentiation we used generalized dissimilarity modelling (GDM). GDM uses I-spline functions to perform non-linear regression between pairwise matrices (Ferrier et al. 2007). The approach has been extensively used to analyze community dissimilarity but has also gained popularity as a way to identify the drivers of genetic differentiation (e.g., Myers et al. 2019; Dool et al. 2022). We performed the analyses on the gdm v1.5.0.1 R package (Fitzpatrick et al. 2021). We specified scaled *â* as our measure of genetic dissimilarity and included geographic distance, taxonomy, and environmental variables as predictors. We classified individuals into the two subspecies of *V*. *tristis* based on their distribution and morphology (when available; Fig 1), coding them as 0 and 1. On the other hand, we extracted 19 bioclimatic variables (Fick and Hijmans 2017) for each point from rasters with 2.5’ resolution. In addition to the present environmental conditions, we also extracted the 19 variables from datasets of past climate obtained from PaleoClim (Brown et al. 2018) corresponding to the Last Interglacial (LIG) (Otto-Bliesner et al. 2006) and Last Glacial Maximum (LGM) (Karger et al. 2021). We also obtained 16 variables for the mid-Pliocene warm period (3.205 Ma) (Hill 2015), as it roughly coincides temporally with the basal split within *V*. *tristis*. We conducted individual analyses for each climatic dataset, specifying as predictors the first principal components of the climatic variables accounting for over 99% of variance. We partitioned the deviance explained by the GDM into the three major groups of variables (i.e., geographic distance, taxonomy, and environmental conditions). Additionally, we used the function “gdm.varImp” to perform significance testing on the model and variables based on 100 rounds of matrix permutation.

### Phylogenetics

We estimated phylogenetic relationships between the individuals and populations of *V*. *tristis* to test for reciprocal monophyly between geographic clusters, which is an indicator of reproductive isolation. Furthermore, phylogenetic hypotheses are required for some of our downstream analyses. First, we inferred the relationships between individuals based on the SNP dataset that included outgroups, totaling 91 individuals and over 50,000 SNPs (Tables S1–S2). We implemented two approaches, concatenated maximum likelihood and a summary method based on the coalescent. We conducted the maximum likelihood analysis in IQ-TREE 1.6.8 (Nguyen et al. 2015). The Bayesian Information Criterion (BIC) was used to compare the fit of several substitution models that account for the ascertainment bias in SNP datasets (Kalyaanamoorthy et al. 2017). The BIC favored the TVM+F+ASC+R3 model.

Support was calculated based on 1,000 ultrafast bootstrap (UFB) replicates (Hoang et al. 2018) and site concordance factors (sCF) (Minh et al. 2020). The coalescent-based analysis was conducted in SVDquartets (Chifman and Kubatko 2014) as implemented in PAUP* 4.0 (Swofford 2003). We obtained bootstrap support (BB) based on 100 pseudo-replicates.

We estimated the relationships between the populations identified by sNMF using SVDquartets. To minimize the impact of introgression, we only included individuals with ancestry coefficients ≥ 0.95 for a given population. We assigned two individuals of *V*. *scalaris* to their own taxon because they did not group with other *V*. *scalaris* in the individual-level analyses. We obtained bootstrap support (BB) based on 100 pseudo-replicates. We time-calibrated the resulting tree with MCMCTree in PAML 4.8 (Yang 2007), specifically using the approach that relies on approximate likelihood to efficiently calculate divergence times for large datasets (dos Reis and Yang 2011). For each population, we kept the individual with the fewest missing data. We specified the concatenated alignment (including invariant sites) as input (262,348 bp). We first estimated branch lengths using BASEML (Yang 2007). Time-calibration relied on a secondary calibration based on the results of Brennan et al. (2021). We specified a uniform distribution with a soft bound on the maximum age. Specifically, we calibrated the split between *V*. *tristis* and *V*. *varius*, with minimum and maximum ages equaling 18 and 24 million years, respectively. The parameters of the distribution were calculated with ‘MCMCtreeR 1.1’ (Puttick 2019). Inference was based on the HKY+G4 substitution model and a global clock model. The Dirichlet-gamma prior for the mean substitution rate was set to *α* = 1 and *β* = 18.7957, based on the mean genetic distance between *V*. *varius* and the members of the *Odatria* subgenus of *Varanus* and the mean age of their most recent common ancestor (Brennan et al. 2021; Pavón-Vázquez et al. 2022). We performed two independent runs with 5,000 burn-in iterations and 2 million post-burn-in iterations sampled every 100 iterations. We combined the runs after verifying convergence and that the effective sample size (ESS) for all parameters was above 200 in Tracer 1.7.2 (Rambaut et al. 2018).

We estimated an explicit network to infer migration between populations. Admixed individuals (i.e., those with ancestry coefficients < 0.95 for any given population) were included in these analyses. *Varanus glauerti* was specified as the outgroup. Analyses were performed using TreeMix 1.13 (Pickrell and Pritchard 2012), a program that takes allele frequency data and detects migration edges (*m*) based on a Gaussian approximation to genetic drift. We specified a block size of 500 SNPs and ran five analyses for each value of *m* between one and five. We used the ‘OptM 0.1.3’ R package (Fitak 2021) to select the optimal number of *m*. This was based on a non-linear least squares model, which had the best fit based on the Akaike Information Criterion (AIC). We verified that the five runs for the best value of *m* were consistent with each other.

Finally, we estimated a Bayesian mitochondrial phylogeny with BEAST 2.7.1 (Bouckaert et al. 2014). We used PartitionFinder (Lanfear et al. 2012) and ModelFinder (Kalyaanamoorthy et al. 2017) to find the optimal partitioning scheme (Chernomor et al. 2016) and corresponding substitution models. The best model included three partitions (best-fitting models in parentheses): the first codon position of *ND4* and *tRNA-His* (K3Pu+F+G4), the second codon position of *ND4* (TN+F+I), and the third codon position of *ND4* (TIM3+F). We specified a coalescent constant population model and a strict clock. We time-calibrated the tree based on a normal prior with a mean of 0.00805 substitutions/site/million years and standard deviation of 0.001 (Bryson et al. 2012; Blair and Bryson 2017). We performed two independent runs, each consisting of 20 million generations with a sampling frequency of 1,000. The runs were combined after assessing for convergence and ESS values above 200 in Tracer. We report the maximum clade credibility tree with 10% burn-in and common ancestor heights.

### Molecular Species Delimitation

We implemented three species delimitation methods using the molecular data: an approach based on unsupervised machine learning without an underlying biological model, an analysis based on the presence of fixed differences, and an approach based on the multi-species coalescent. Namely, the first method is based on self-organizing maps (SOM) (Kohonen 1998; Wehrens and Buydens 2007) and was recently used for species delimitation in a taxonomically challenging group (Pyron et al. 2023). SOM is an artificial neural network that reduces dimensionality based on similarity (Wehrens and Buydens 2007). Briefly, an input vector is projected into a two-dimensional output grid based on clustering in multi-dimensional space. Adjacency between cells in the output grid indicate similarity. After classifying observations into grid cells, *k*-means clustering can be used to identify the optimal number of clusters (i.e., species, in this context) (Wehrens and Buydens 2007; Pyron et al. 2023). Analyses were performed in the ‘kohonen 3.0.10’ R package (Wehrens and Kruisselbrink 2018). We used the same data as for sNMF to calculate allele frequencies for each individual to be used as input. We specified an output grid with 9 x 9 cells, so that each individual could potentially be classified into its own cell. We performed 50 replicates of the classifying algorithm, each consisting of 400 iterations and with a learning rate *α* (0.05, 0.01). Using *k*-means clustering, we selected the optimal number of clusters based on the maximum sequential decrease in the weighted sum of squares across all runs.

Our second approach is the fixed difference analysis (FDA) introduced by Georges et al. (2018). Fixed allelic differences between populations are strong indicators of reproductive isolation (Kozak and Montanucci 2001; Georges et al. 2018). We tested for fixed differences between the populations identified by sNMF using the “gl.fixed.diff” function of ‘dartR’. The approach relies on simulation to account for the probability of false positives arising because of the incomplete sampling of populations. We specified 1,000 simulation iterations and considered differences as “fixed” if one allele is present in ≥ 95% of the individuals in one population and ≤ 5% of individuals in the other. We then repeated the analysis after collapsing the populations that did not show significant fixed differences in the first round.

Our third approach is based on the genealogical divergence index (GDI) (Jackson et al. 2017; Leaché et al. 2019). The GDI of taxon A equals 1 – *e* ^−2*τ*AB/^ *^θ^*^A^, where *τ*AB is the divergence time between taxa A and B, and *θ*A is the effective population size of A. Panmixia is indicated by GDI = 0, whereas GDI = 1 indicates complete isolation. Values below 0.2 are usually considered to indicate con-specificity, values between 0.2 and 0.7 are ambiguous, and values above 0.7 are considered to be strong evidence for species-level divergence (Jackson et al. 2017; Leaché et al. 2019). We used BPP 4.1.4 (Flouri et al. 2018) to estimate *θ* and *τ*, and used these estimates to get a posterior distribution of the GDI. We followed the approach of Leaché et al. (2019), where pairs of sister taxa are recursively collapsed based on their GDI. Given computational constraints, we performed the analysis on a random sample of 500 loci and 28 individuals selected to maximize the representation of genetic variation. The sequences were phased using fastPhase 1.4.8 (Scheet and Stephens 2006). We estimated parameters on the fixed population tree estimated by SVDquartets. The inverse-gamma distribution for the priors of *θ* and *τ* were based on the mean genetic distance within populations and the mean distance between the earliest diverging population and the rest, respectively. For each round of testing, we conducted two analyses consisting of 100,000 burn-in iterations and 2 million post-burn-in iterations sampled every second iteration. The runs were combined after assessing convergence and verifying that the ESS for each parameter was above 200 in Tracer.

### Morphometric Analyses

We obtained four morphometric datasets to test whether two putative species proposed by the molecular analyses are morphologically divergent: snout-vent length (SVL; used as proxy for body size); 17 linear measurements describing body shape; a two-dimensional geometric morphometric dataset describing head shape in dorsal view, consisting of 13 landmarks and 20 sliding semi-landmarks; and a two-dimensional geometric morphometric dataset describing head shape in lateral view, consisting of 10 landmarks. All the datasets were recorded by the first author after examining specimens deposited across seven herpetological collections (Tables S3–S5). Our sampling includes the types of *Odatria punctata* Gray 1838 and *Varanus punctatus* var. *orientalis* Fry 1913. We processed the data following Pavón-Vázquez et al. (2022). Briefly, missing measurements (i.e., from incomplete tails) were imputed using random forest training (Stekhoven and Bühlmann 2012). SVL was log-transformed, while log-shape ratios were calculated for the body shape measurements (Mosimann 1970). We used generalized Procrustes analysis (GPA) (Gower 1975) in ‘geomorph 4.0.0’ (Adams et al. 2022; Baken et al. 2021) to standardize the geometric morphometric datasets. For each dataset, we used the “procD.lm” function of ‘geomorph’ to test for sexual dimorphism. For a dataset exhibiting sexual dimorphism (body shape), we deleted females because our sampling is male-biased. Our final datasets included 62 (SVL, dorsal head shape), 45 (body shape), and 61 (lateral head shape) adults of *V*. *tristis*.

We implemented analyses of variance to detect significant morphometric differences between the putative species. Those individuals that were sequenced were assigned to putative species as indicated by the molecular data. Other individuals were classified based on their geographic origin with respect to the nuclear clusters. For each linear measurement, namely SVL and the 17 body measurements, we used R to perform an ANOVA. For the multivariate datasets, including the body shape dataset, we used the “procD.lm” function of ‘geomorph’, relying on 10,000 iterations of residual randomization permutation (Collyer et al. 2015) for significance testing. To visualize the differences between the groups, we used boxplots for SVL and PCA for the multivariate datasets.

### Coloration Analyses

We tested whether geographic variation in head color is indicative of gene flow or isolation between the putative species. Our analyses are based on the examined specimens and photographic records issued by the wider public. We downloaded the photographic records using ‘galah 1.5.1’ (Westgate et al. 2023), an R interface to the network of biodiversity living atlases. In our case, we downloaded the photographs and their associated metadata from the Atlas of Living Australia (Belbin et al. 2021). We did not include photographs with missing coordinates, where head coloration could not be clearly distinguished, and those of juveniles. The first author classified individuals into the following head color categories: “black”, when any pattern (cephalic reticulation and dark post-ocular stripes) is mostly obscured by black pigment; “dim”, when some patterning is visible, but the head is still noticeably darker than the rest of the body; “pale”, when the head is light and the pattern is faded; “yellow”, when the head is yellowish and the pattern is faded; and “freckled”, when the head is light and the patterning is clearly visible on the head and neck. A map following this classification is shown in Fig. S1. However, in our analyses we collapsed the “black” and “dim” categories into a single “dark” category, and the “pale” and “yellow” categories into a single “light” category. There were two reasons for this: one was to increase sample sizes and the second was that it can be hard to tell apart the subcategories within the collapsed categories in preserved specimens. We included adults of both sexes in our analyses, after verifying that any given sex could show any of the patterns and that variation was basically geographical.

Mapping revealed that the “freckled” pattern is exclusively found in one of the putative species. Our analyses were thus aimed at testing whether variation in the other putative species is indicative of gene flow with the former or is instead reflecting local adaptation or plasticity within the latter. To test this, we fitted Bayesian generalized nonlinear regression models in the ‘brms 2.19.0’ R package (Bürkner 2017), implementing a modified version of the approach described by Hantak et al. (2022). Color was indicated as a response variable under a Bernoulli distribution; i.e., we modelled the probability of being light-colored. One of the predictors was the minimum geographic distance to freckled individuals. If light color is the result of ongoing gene flow with the freckled form, we anticipated that the probability of being light would increase with proximity to freckled individuals. We measured geographic distance as great-circle distance using the ‘codep 0.9.1’ R package (Guenard et al. 2018). To test whether variation is environmentally driven instead, we included the following predictors: latitude, elevation, annual mean solar radiation (averaged across months), and annual precipitation. Environmental rasters had 30” resolution and were obtained from WorldClim2 (Fick and Hijmans 2017). Latitude was multiplied by –1 and all predictors were log-transformed and scaled. We also obtained annual mean temperature for each individual but did not include it in the models because it was tightly correlated (*R*^2^ ≥ 0.40) with other predictors. After deleting pseudo-replicates—individuals that were in the same raster cells as others—we retained 149 individuals.

We fitted a total of 31 models specifying every possible combination of predictors. For each model, we ran four chains with 1,000 burn-in iterations and 2,000 post-burn-in iterations. We compared model fit using the leave-one-out cross-validation criterion (Vehtari et al. 2017). To evaluate how much variation is accounted for by the models we calculated the Bayesian version of *R*^2^ with the “bayes_R2” function. We plotted the conditional effects of the optimal model after verifying that adding interaction terms did not increase fit and that there was no significant spatial autocorrelation based on Moran’s *I* using the ‘pgirmess 1.6.9’ R package (Giraudoux 2018).

### Integrative Species Delimitation

We used the SuperSOM approach to infer species limits based on the simultaneous analysis of multiple types of data. Unlike the SOM analyses presented above for the molecular data, SuperSOM can reduce dimensionality based on multiple input layers (Wehrens and Buydens 2007; Pyron 2023). Weighing of the layers is independent of scale or the amount of data in each layer.

Our SuperSOM analysis was based on molecular, phenotypic, spatial, and environmental data. We implemented the approach of Pyron (2023) and used the scripts accompanying that publication, which rely primarily on the ‘kohonen’ R package. Since the effects of missing data are poorly known (Pyron 2023), we conducted this analysis on 23 individuals for which all types of data are available. This reduced sampling still sufficiently covers the geographic range of *V*. *tristis*. The *V*. *tristis* DArTseq dataset was processed to ensure reproducibility and informativeness for this smaller sample (Table S2). The spatial data included latitude, longitude, and elevation. The environmental data included mean solar radiation and climatic variables representing annual averages, extremes, and seasonality (bioclimatic variables 1, 4– 6, and 12–15) at 30” resolution (Fick and Hijmans 2017). The phenotypic matrix included log-scaled SVL, the log-shape ratios of the 17 linear measurements, the Procrustes-aligned coordinates of the head shape datasets (aligned anew for these set of individuals), and color (coded as 0 = dark, 0.5 = light, and 1 = freckled). All the non-genetic variables were scaled through min-max normalization. We specified 4 x 4 cells for the output grid and performed 100 replicate analyses, each consisting of 100 iterations with a learning rate *α* (0.5, 0.1). We follow Pyron (2023) and use the term “species coefficients” to denote the frequency of assignment of each individual to a particular cluster across the replicates.

## Results

### Population Structure

The sNMF analyses recovered six as the optimal value of *K* based on cross-entropy. The six populations are as follows (Fig. 3a; Table S6): a widespread population (WS), with a wide distribution in the Arid Zone; a southwest population (SW), from the Mediterranean areas in southwest Australia; a central population (CT), confined to the Musgrave Ranges; a population found in the Top End (TE) in northern Australia; a southeastern population (SE) with a wide distribution within the Great Artesian Basin; and a northeastern population (NE), occurring east of the northern portions of the Great Dividing Range. WS shows signs of substantial admixture with each of the populations where they approach each other, except for NE. Substantial admixture of WS with SE is restricted to a single individual from Lake Buchanan, Queensland. We did not find signs of admixture between NE and SE where they approach each other, even when our sampling includes a region of possible sympatry around Lake Buchanan. The PCA mirrored these results (Fig. 3b). Scores for the first principal component, accounting for 26.46% of variance, differentiate NE from other populations. SE groups separately from the remaining populations along the second principal component, accounting for 8.49% of variance. The Lake Buchanan individual is roughly halfway between the latter two clusters. The scores for the third principal component differentiate TE (Fig. S2).

**FIGURE 3.**
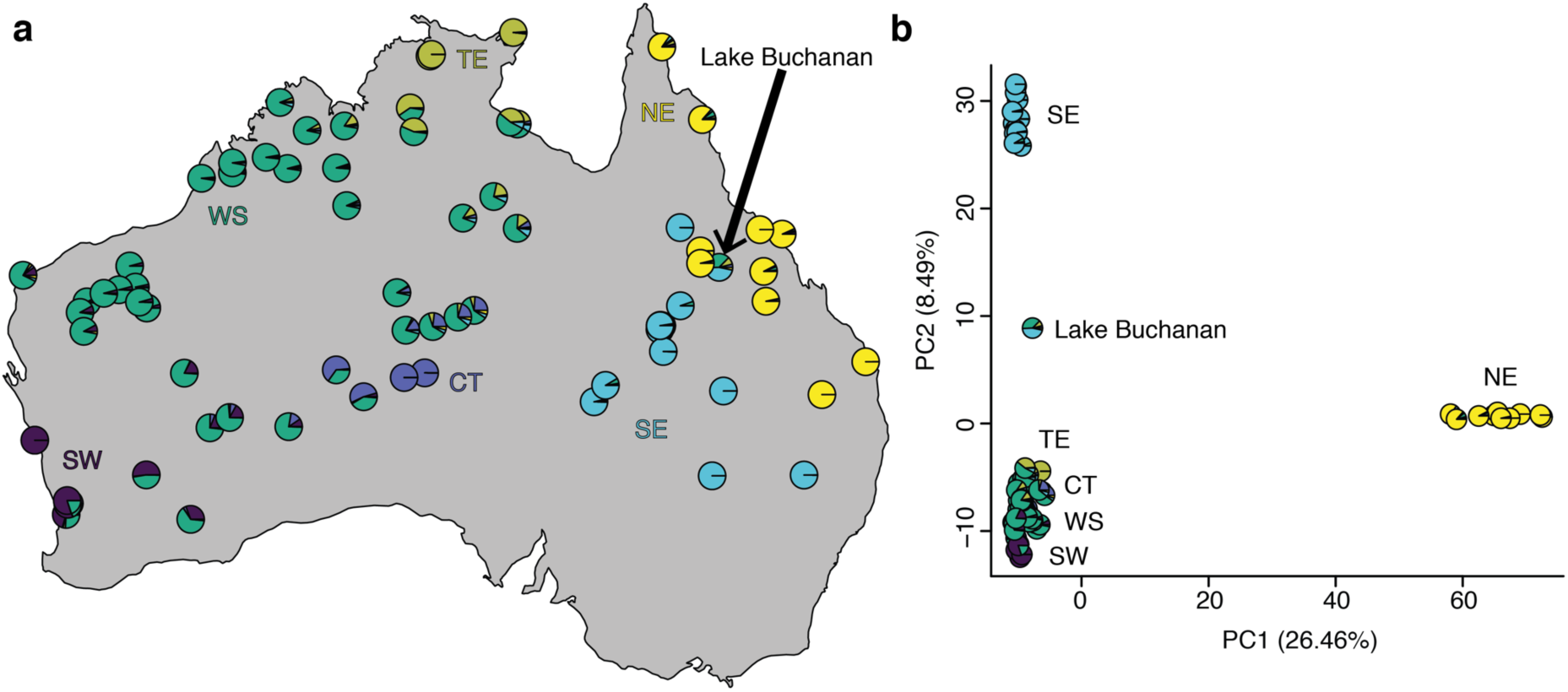
Population structure in *Varanus tristis*. a) Ancestry coefficients estimated with sNMF for *K* = 6; each pie chart represents an individual. b) Principal component analysis of SNP data; pie charts indicate sNMF ancestry coefficients for individuals.

### Patterns of Isolation

The Mantel test revealed a significant positive relationship between genetic and geographic distance (*r* = 0.29; *p* = 0.001). However, visual inspection revealed that for similar geographic distances the genetic distances between NE and other populations are disproportionately high (Fig. 4a). Similarly, effective migration rates (Fig. 4b) are particularly low in the geographic area separating NE from other populations. This region corresponds to the border between the Great Dividing Range and Great Artesian Basin. Other areas with low effective migration are found in the borders between the populations identified by sNMF.

**FIGURE 4.**
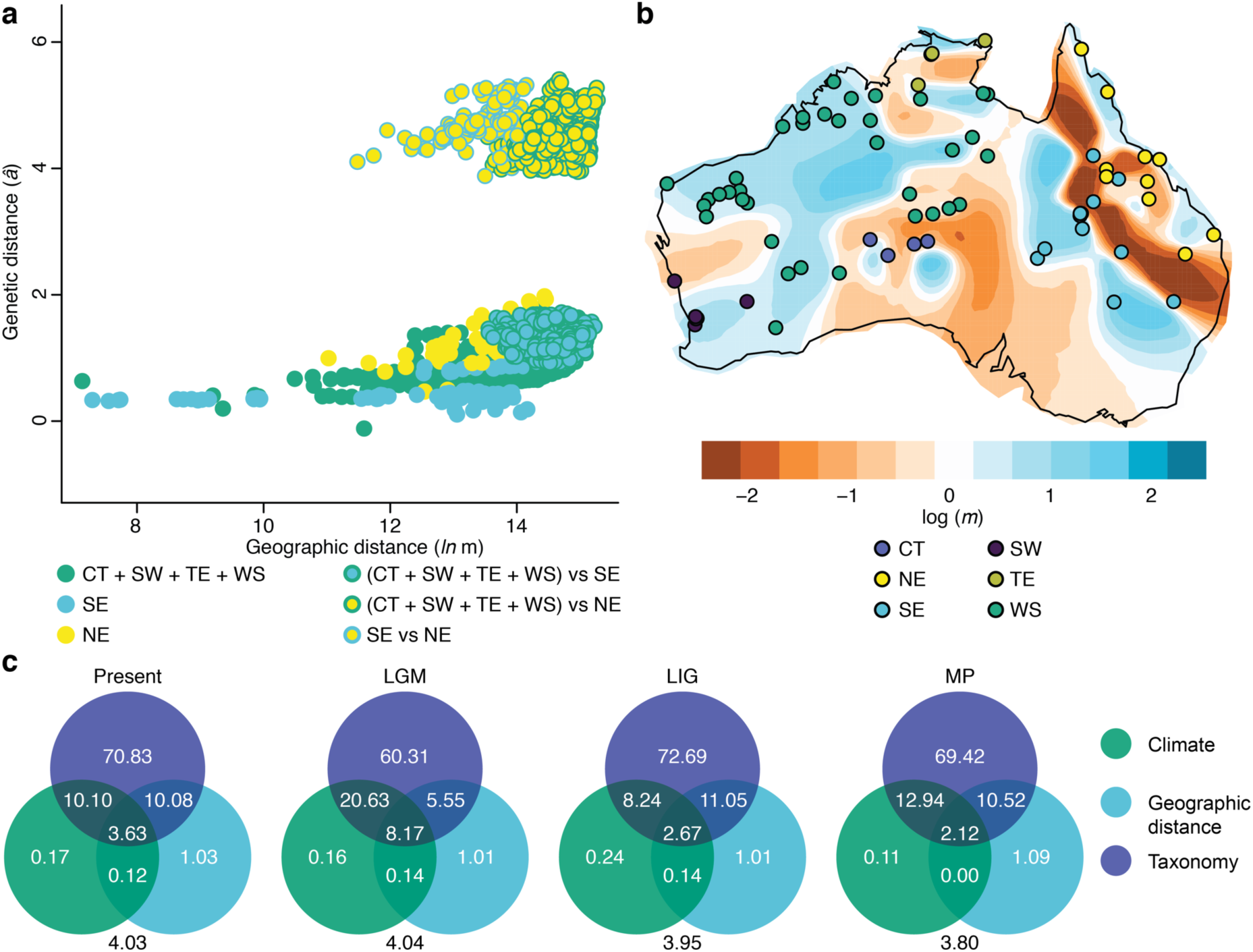
Patterns of isolation in *Varanus tristis*. a) Relationship between genetic and geographic distance; each point represents a pairwise comparison between individuals; comparisons are colored to allow comparison between three major clusters identifiable by their principal component scores (Fig. 3b). b) Log-scaled effective migration rates (*m*); positive values indicate that *m* is larger than expected under isolation by distance, negative values indicate it is smaller; individuals are colored by population. c) Deviance in genetic distances explained by climate, geographic distance, and taxonomy (i.e., whether individuals are assignable to *V*. *t*. *tristis* or *V*. *t*. *orientalis*) obtained through generalized dissimilarity modelling; climatic variables were extracted from four points in time: the present, Last Glacial Maximum (LGM), Last Interglacial (LIG), and mid-Pliocene warm period (MP); values at the center of the circles show the deviance explained by each set of variables alone, values at the intersections are the deviance explained by that section of the intersection alone (i.e., the sum of the values within a circle give the total deviance explained by each set of variables).

The GDM (Fig. 4c; Tables S7–S9) indicated that taxonomy is a strong predictor of genetic disparity. Across all the time slices, the assignation to subspecies explained the largest proportion of the deviance in genetic distances, both independently and jointly with other variables. When taken independently, geographic distance explains a larger proportion of the deviance in genetic distances than climate. However, their joint contributions are comparable. Climate during the LGM explains a particularly large proportion of the deviance in genetic distances compared to other points in time. The full models were significant across all time slices based on matrix permutations.

### Phylogenetics

The individual-level trees obtained with SVDquartets (Figs. 5a, S3) and IQ-TREE (Fig. S3) were similar to each other. In both, *V*. *scalaris* was not monophyletic. Two samples of *V*. *scalaris* from the Wet Tropics of northeastern Queensland (hereafter *V*. *scalaris* WT) were recovered as sister to a clade containing *V*. *glauerti* and *V*. *tristis* with strong support. The reciprocal monophyly of the latter two taxa was also strongly supported. Samples assigned to NE conform a strongly supported clade that is sister to the rest of the sampled *V*. *tristis*. WS was not monophyletic. Instead, moderately to strongly supported clades corresponding to CT, SE, SW, and TE are nested within WS.

**FIGURE 5.**
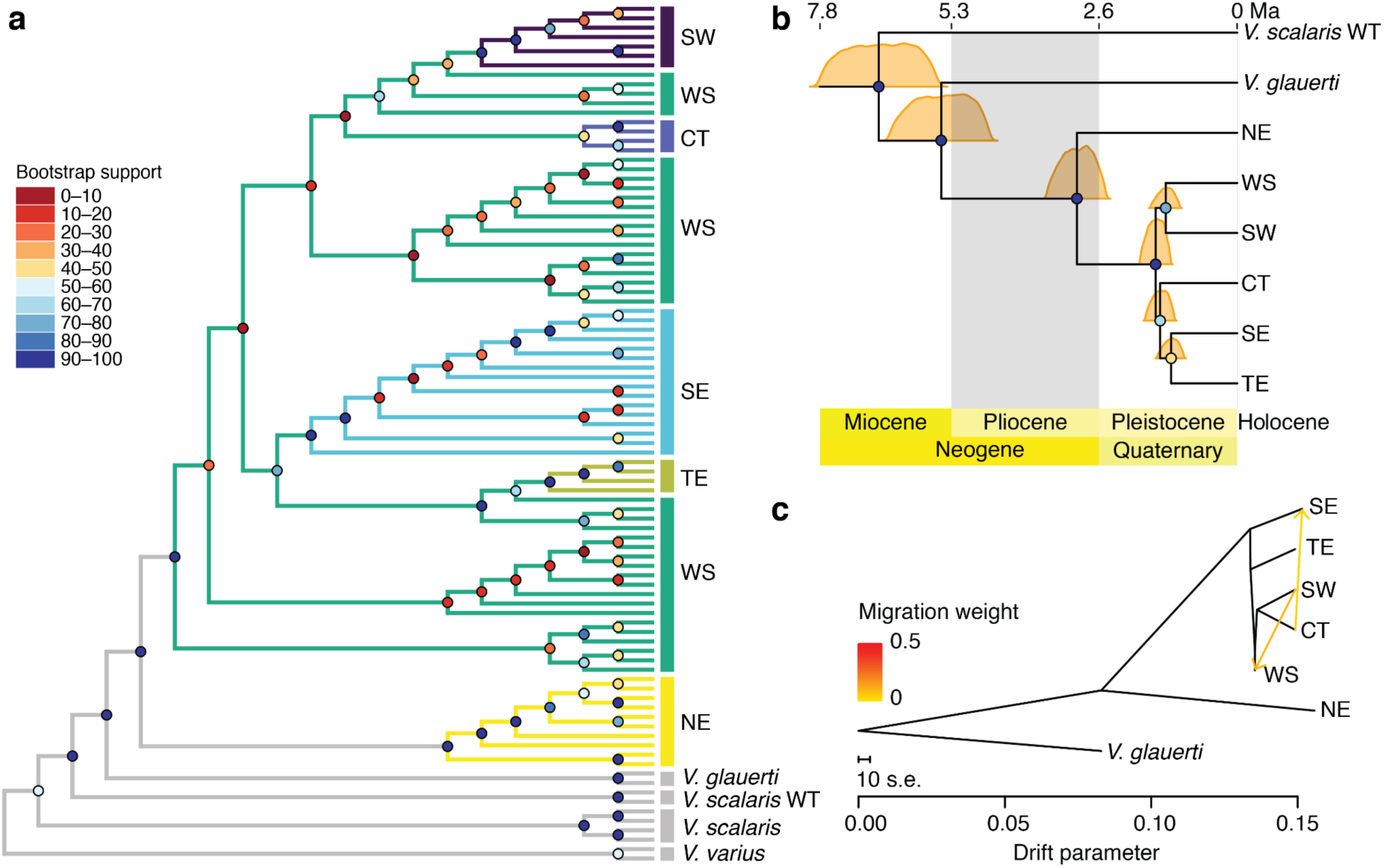
Phylogenomics of *Varanus tristis*. a) Individual-level phylogeny based on the SNP data and estimated with SVDquartets; circles at nodes indicate bootstrap support; branch lengths are arbitrary. b) Time-calibrated, population-level tree estimated with SVDquartets and calibrated with MCMCTree; posterior distributions of divergence times are shown for each node. c) Population network obtained with TreeMix under the optimal number of migration edges (two); branch lengths are proportional to estimated genetic drift; the smaller scale bar indicates ten times the average standard error of the sample covariance matrix.

The population-level tree estimated with SVDquartets (Fig. 5b) also recovered *V*. *scalaris* WT as sister to *V*. *glauerti* + *V*. *tristis*, to the exclusion of other *V*. *scalaris*. Within *V*. *tristis*, NE was recovered as sister to the rest of the populations. The clade comprising the latter populations was supported strongly, but the relationships between them were not. SW and WS were recovered as sister, whereas CT appeared as sister to SE + TE. Dating suggests that the split between *V*. *glauerti* and *V*. *tristis* happened during the late Miocene (posterior mean [PM] = 5.52 Ma; 95% highest posterior density [HPD] = 4.63–6.48). The basal split within *V*. *tristis* likely happened during the Late Pliocene (PM = 2.99 Ma; 95% HPD = 2.46–3.55). Other divergences within the taxon were dated to the Pleistocene.

TreeMix (Fig. 5C) also recovered NE as sister to a clade conformed by the rest of the populations. The topology recovered for the latter was (SE, (TE, (WS, (CT, SW)))). The non-linear least squares model suggested that the optimal value of *m* was two (Fig. S4). Migration was recovered from SW into WS and from CT into SE.

The mitochondrial tree (Fig. S5) recovered *V*. *scalaris* WT as sister to topotypic *V*. *glauerti*, to the exclusion of other *V*. *scalaris*. While the sister relationship between *V*. *scalaris* WT and *V*. *glauerti* was not supported strongly, a clade comprising *V*. *scalaris* WT, *V*. *glauerti*, and *V*. *tristis* was. A sample of *V*. *glauerti* from Arnhem Land in the Top End was nested within *V*. *tristis*. That sample could not be included in the SNP analyses because the tissue sample was depleted and we could not locate any DNA extractions. Samples assignable to NE formed a strongly supported clade that was sister to the rest of *V*. *tristis* plus the *V*. *glauerti* sample from the Top End. The latter sample appeared as sister to the rest of *V*. *tristis*, but its position was poorly supported. The remaining samples of *V*. *tristis* conformed clades that roughly correspond geographically to the populations recovered by sNMF. However, the sNMF clusters did not form reciprocally monophyletic groups. Divergence dates were generally older than those estimated with the genomic data. The split between NE and its sister group was inferred to be 4.23 Ma old (95% HPD = 3.30–5.29). The most recent common ancestor of the remaining *V*. *tristis* was estimated to be 2.91 Ma old (95% HPD = 2.20–3.67).

### Molecular Species Delimitation

Forty-one of the cells in the 9 x 9 SOM output grid were occupied by 1–4 individuals each (Figs. 6a). Division into two clusters was supported by *k*-means clustering, since this value of *k* showed largest decrease in weighted sum of squares (both absolute and averaged) across 50 independent replicates (Fig. S6). The resulting clusters corresponded to NE and the rest of the populations together. The assignment probability of all individuals to their corresponding clusters was one, based on the 50 replicates.

**FIGURE 6.**
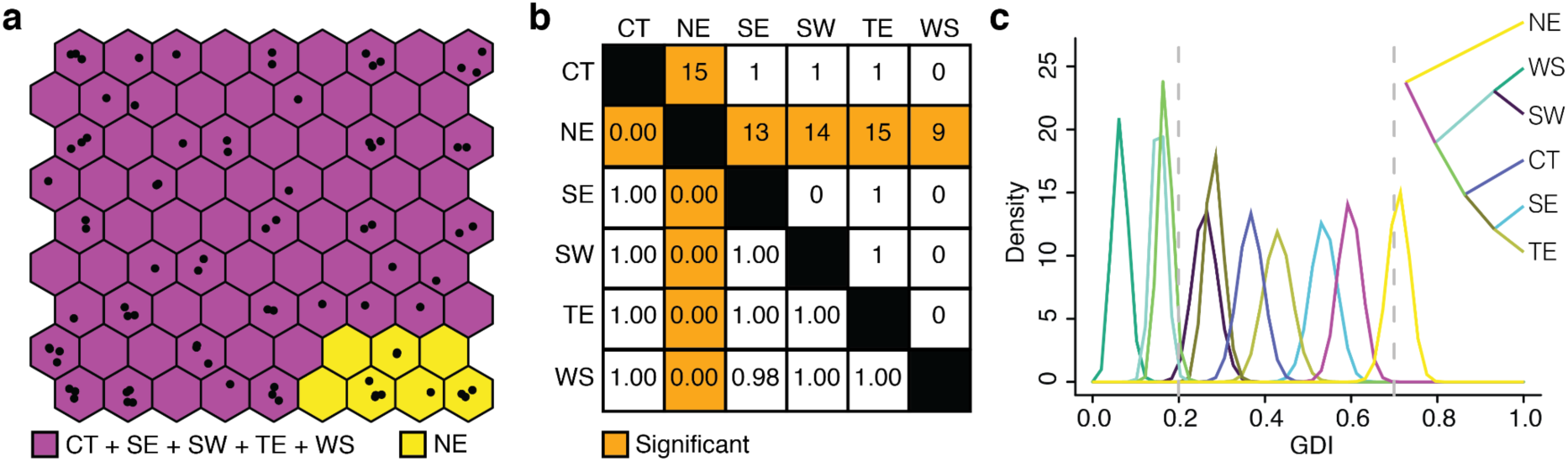
Species limits in *Varanus tristis* based on molecular data. a) Output grid obtained through self-organizing maps; cell adjacency indicates similarity in allele frequencies; cells are colored according to their assignment to groups based on *k*-means clustering; each point corresponds to an individual. b) Fixed difference analysis; the upper triangle indicates the percentage of fixed allelic differences between populations; the lower triangle indicates *p*-values based on simulations to account for false positives arising from sampling error; significant comparisons are highlighted. c) Posterior distribution of the genealogical divergence index (GDI) for each branch in the SVDquartets population-level tree; sister terminals were recursively collapsed based on their GDI; GDI < 0.2 suggests con-specificity, 0.2 < GDI > 0.7 indicates ambiguous results, and GDI > 0.7 suggests species-level divergence.

The first round of FDA (Fig. 6b) detected significant fixed differences between NE and every other population. There were ≥ 9% fixed allelic differences between the former population and the others. After collapsing the remaining populations into a single unit, we again found significant differences with NE (*p* < 0.001; fixed allelic differences = 8%).

Regarding the GDI analyses (Fig. 6c), NE had the highest GDI. The mean and most of the posterior distribution are above the 0.7 threshold, indicating strong evidence for speciation. Most of the other branches showed ambiguous values of GDI (0.2 < GDI > 0.7). Within this category, SE and the cluster comprising all the populations except for NE had the highest GDI. Given the above evidence, we treated NE (corresponding to *V*. *t. orientalis*) and the remaining populations (corresponding to *V*. *t*. *tristis*) as putative species in the morphological analyses.

### Morphometric Analyses

We found that while individuals of *V*. *t. tristis* are generally larger than those of *V*. *t*. *orientalis* (Fig. 7a), their mean SVLs were not significantly different (*F* = 3.16; *p* = 0.0806). Regarding our linear measurements describing body shape, we found significant differences in the length of the fourth finger (*F* = 9.48; *p* = 0.0036), hand width (*F* = 4.10; *p* = 0.0490), length of the fourth toe (*F* = 6.98; *p* = 0.0114), and foot width (*F* = 14.07; *p* = 0.0005). The multivariate analyses indicated significant differences in body shape (*F* = 2.61; *p* = 0.0065) (Fig. 7b), dorsal head shape (*F* = 2.55; *p* = 0.0380) (Fig. 7c), and lateral head shape (*F* = 2.45; *p* = 0.0111) (Fig. 7d). Other results are summarized in Table S10.

**FIGURE 7.**
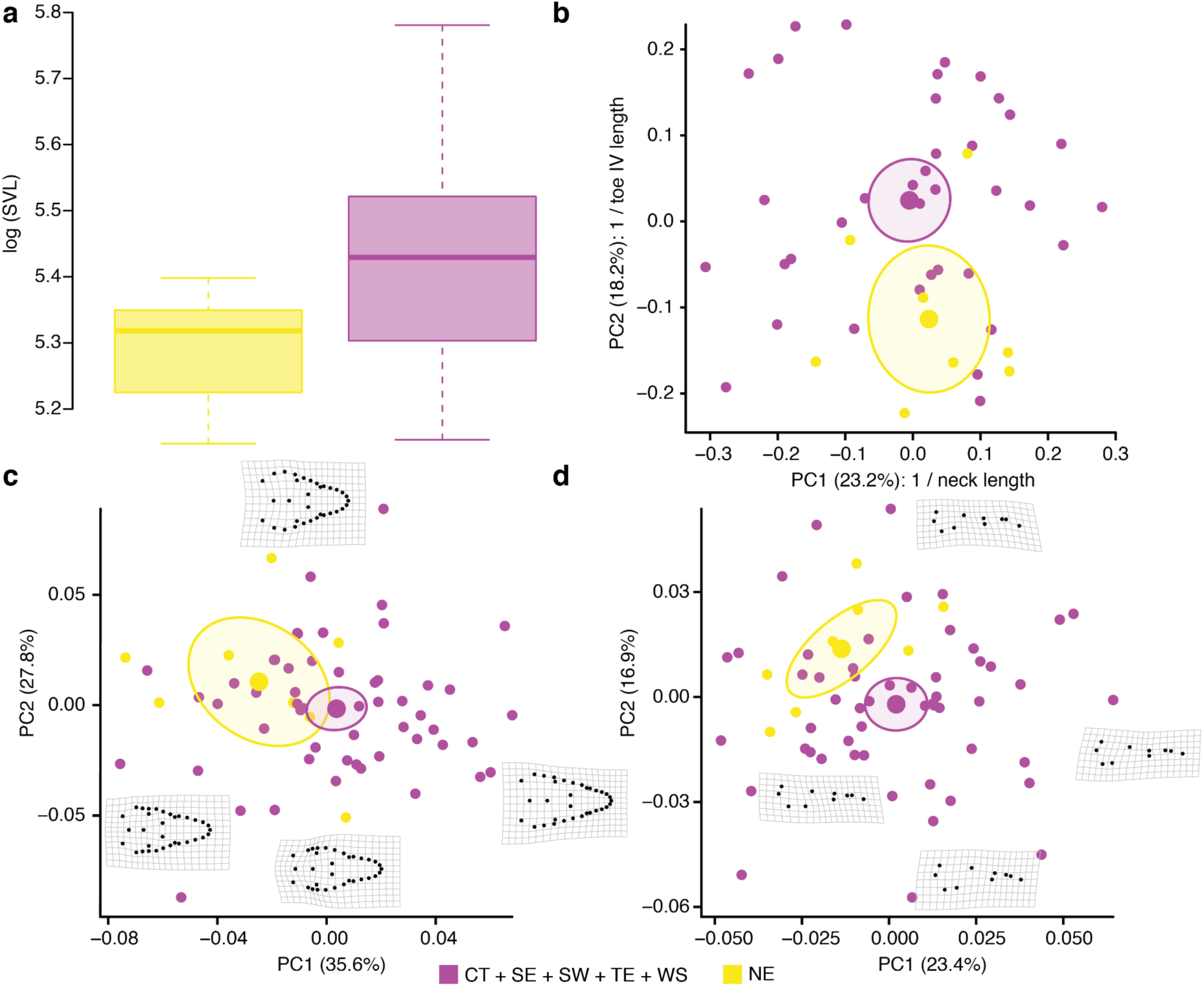
Morphometric variation in *Varanus tristis*. a) Boxplot of log-transformed snout-vent length (SVL). b) Plot of principal component analysis (PCA) for body shape; small points represent individuals, large points represent the mean for each putative species, circles indicate 95% confidence ellipses; the variables contributing the most to variation in each PC axis are shown. c) PCA plot for head shape in dorsal view; for the individuals at the extremes of each PC axis, the landmark configuration and deformation grid with respect to average shape are shown. d) PCA plot for head shape in lateral view; same as for Fig. 7c.

### Coloration Analyses

Color variation in *V*. *tristis* (Fig. 8a) appears to be geographically structured. We found that the “freckled” pattern is found exclusively in northeastern Australia (Fig. 8b), corresponding to the distribution of *V*. *t. orientalis*. Variation in *V*. *t*. *tristis* appears to follow a latitudinal gradient (Fig. 8b). Light colored individuals are mostly found in the northern portion of the distribution. This was corroborated by our Bayesian models. The best-fitting model according to the leave-one-out cross-validation criterion was that including latitude and solar radiation as predictors (Table S11). The PM of *R*^2^ for this model is 0.65. The model suggests that there is a higher probability of observing light colored individuals at higher latitudes and where solar radiation is less intense (Fig. 8c). The best-fitting model based on a single variable was that including latitude (expected log pointwise predictive density [elpd] difference with best model = 0.7; PM *R*^2^ = 0.63). The model based exclusively on the geographic distance between the putative species performed poorly (elpd difference = –45.6; PM *R*^2^ = 0.04). In fact, it had the worst fit according to the leave-one-out cross-validation criterion.

**FIGURE 8.**
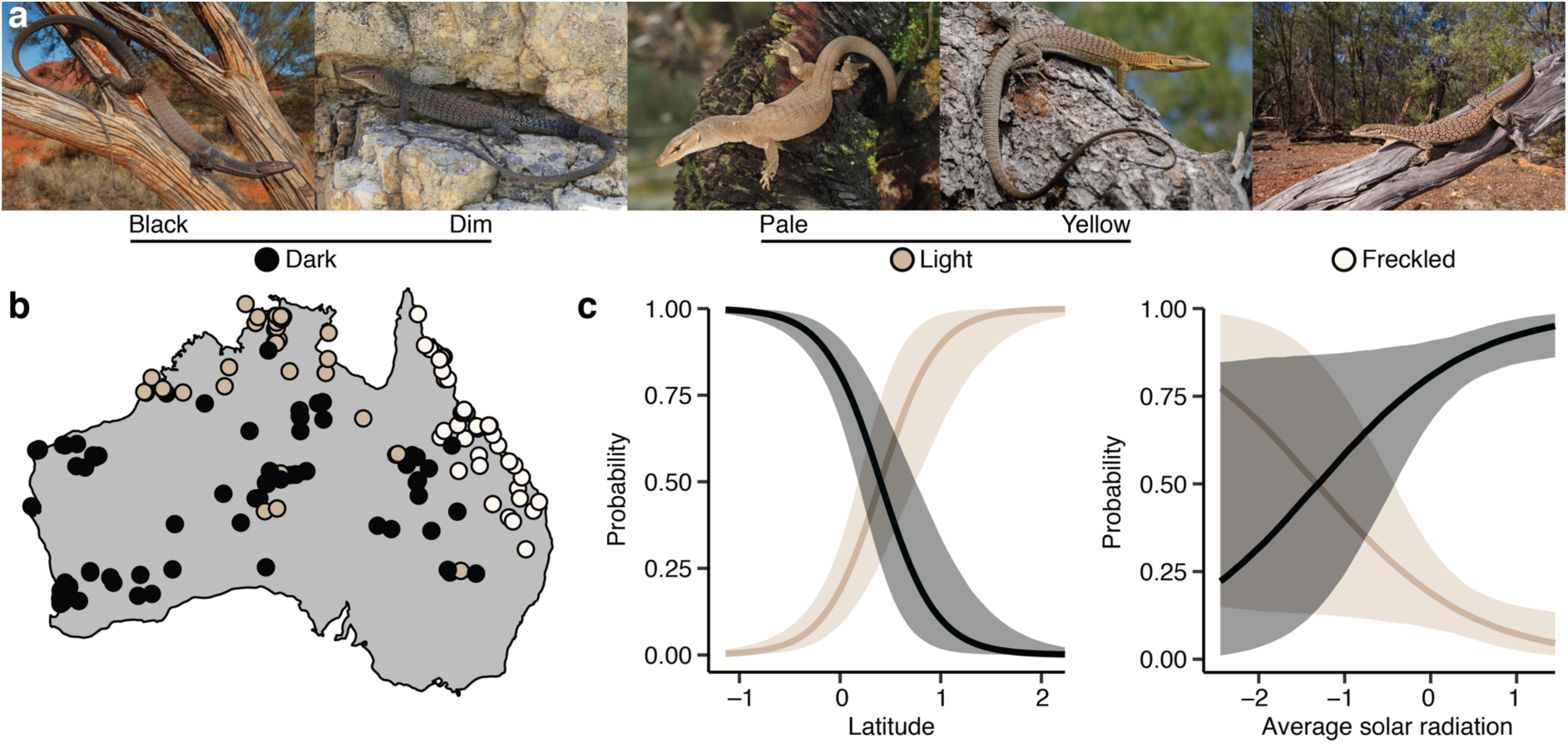
Color variation in *Varanus tristis*. a) Classification of variation into discrete categories; some categories were collapsed to increase sample sizes and because they were difficult to differentiate in preserved specimens; photographs by (left to right) Jules Farquhar, William Scott, Ryan Ellis, Jules Farquhar, and Stephen Zozaya. b) Geographic distribution of the three main color categories. c) Conditional effects under the best-fitting non-linear model; the probability of being light or not was modelled, the probability of being dark was extrapolated from the former; shaded contours indicate 95% credible intervals; the predictors were log-transformed and scaled; geographic distance of dark and light individuals to freckled individuals was included in some models, but these were not favored.

### Integrative Species Delimitation

In our SuperSOM analysis the molecular data had the largest scaled weight, followed by the phenotypic, climatic, and spatial data, in that order (Fig. 9a). Results support two clusters, corresponding to NE on one hand and the rest of the populations on the other (Figs. 9b, S6). Within the 4 x 4 output grid, 13 cells were occupied by 1–3 individuals each (Fig. 9b).

**FIGURE 9.**
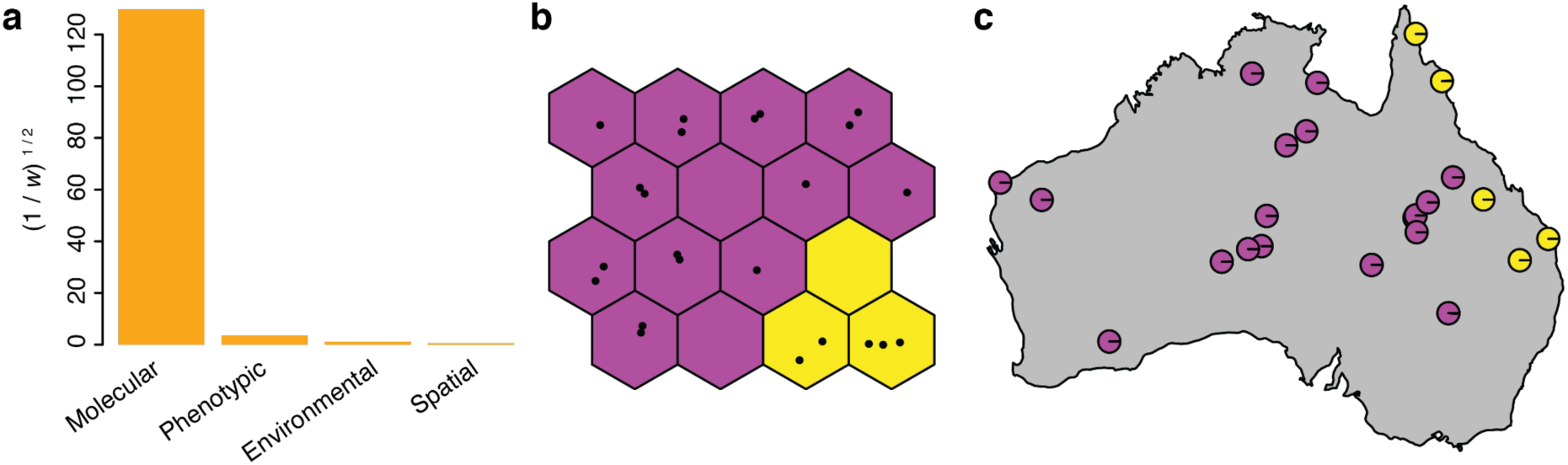
Integrative species delimitation in *Varanus tristis* using multilayered self-organizing maps. a) Scaled relative weights describing the contribution of different types of data. b) Output grid for *k* = 2; as in Fig. 6a. c) Frequency of classification of each individual into different clusters (i.e., species coefficients) obtained through 100 replicates.

Species coefficients for most individuals equaled one (Fig. 9c). For two individuals, a conflicting assignment was recovered by one run each. These individuals come from Gundabooka National Park, north-central New South Wales, and Lambert Creek, northeastern Queensland, respectively.

## Discussion

### Future Directions for Species Delimitation

Many of the components in our workflow are characterized by the delineation of clear hypothesis and expectations (Fig. 2). We also evaluated alternative explanations and accounted for confounding factors when exploring the observed patterns of genetic structure (IBD, IBE, incomplete lineage sorting, gene flow) and phenotypic variation (local adaptation/plasticity). We think these should be important considerations in species delimitation studies to avoid the taxonomic mischaracterization of diversity. Furthermore, they reinforce the status of species as hypotheses (de Queiroz 2005), providing clearer guidelines for other researchers to further test the results of species delimitation studies.

Delimiting species will invariably entail some degree of arbitrariness, as discrete criteria and thresholds have to be applied to a continuous and fractal pattern of divergence (Zachos 2016). However, clearly specifying the criteria used to separate taxonomic entities makes it easier for researchers and decision-makers to evaluate whether it is appropriate for them to incorporate taxonomic proposals from a practical point of view (e.g., whether to treat lineages as distinct units for macroevolutionary studies or conservation). Furthermore, studies that focus on evolutionary processes for species delimitation will produce valuable information about the biology of the organisms at hand and evolutionary processes in general, independently of their taxonomic conclusions.

As demonstrated here, community science has the potential to be a major tool in species delimitation. Records generated by the public can far outnumber those available in collections and better represent species ranges (Chowdhury et al. 2023a). Community observations can also harbor temporal, spatial, ecological, or even behavioral and acoustic information. Thus, they have the potential to shed light on several aspects that are relevant for species delimitation such as variation in mate recognition traits, temporal divergence in breeding activity, or phenotypes at contact zones. However, there are limitations to the utility of community observations posted in public platforms such as iNaturalist. The precise location of threatened taxa is often obscured, most observations are restricted to the Global North (Rosa et al. 2022), and the processing of a large number of records can pose practical challenges. However, these limitations can be ameliorated by putting procedures in place to facilitate access to obscured metadata by researchers, using observations posted on social media (Chowdhury et al. 2023a), and by the implementation of automated image processing workflows (de Solan 2020; Hantak et al. 2022) (see below), respectively.

Artificial intelligence is rapidly evolving as a field and it has the potential to greatly accelerate the discovery and description of biodiversity. Artificial intelligence has the potential to streamline the procurement of large phenotypic datasets in the contexts of both museum collections and community science. Machine learning can be used to quickly gather geometric morphometric data for many specimens with reduced error (Devine et al. 2020).

This kind of algorithms can also be used to extract phenotypic data from photographs obtained under different conditions, as is the case of community observations (de Solan 2020; Hantak et al. 2022). Machine learning can then be used to analyze these data solely or jointly with molecular data to assign them to known species with great accuracy (Wäldchen and Mäder 2018; Valan et al. 2019; Yang et al. 2022). Acoustic data can also be used to efficiently identify species in a deep learning framework (Kahl et al. 2021). This kind of methods can then be extended to recognize variation that falls outside that expected for recognized taxa, i.e., identify undescribed species (Yang et al. 2022).

Finally, machine learning can be used to delimit species outside the context of new species discovery. We implemented some of those approaches here (SOM and SuperSOM). They are promising in that they appear to be conservative. In *V*. *tristis*, they were able to distinguish between population structure and species-level divergence, recovering fewer entities than sNMF. A similar behavior has been reported for other unsupervised machine learning approaches used in species delimitation (Derkarabetian et al. 2019). While these methods are not based on biological models, there are other approaches that rely on data simulated under a variety of evolutionary scenarios for training (e.g., Pei et al. 2018; Smith and Carstens 2020). These workflows mainly employ molecular data because the effects that evolutionary processes have on DNA have been relatively well studied. However, it is also possible to build models that can distinguish between intra and interspecific phenotypic variation (Solís-Lemus et al. 2015). The utility of these models would be amplified if they are expanded to account for a greater variety of evolutionary scenarios such as secondary contact and the reinforcement of reproductive isolation.

### Speciation, Structure, and Variation in V. tristis

Our analyses revealed notable genetic structure and geographic variation in *V*. *tristis*, some likely representing species-level divergence. NE was consistently recovered as the most divergent population. Admixture with other populations was minimal (Fig. 3), its genetic differentiation cannot be explained solely by IBD or IBE (Fig. 4), it is sister to the rest of *V*. *tristis* in both the mitochondrial and nuclear phylogenies (Figs. 5, S3, S5), and the molecular and integrative species delimitation analyses (Figs. 6, 9) support its specific status.

Phenotypic differences were found in the morphometric data (Fig. 7) and more evidently coloration (Fig. 8). Specimens assigned to NE can be distinguished by possessing a distinct dark post-ocular stripe and a reticulated dorsal pattern that extends anteriorly into the head. Geographic proximity with NE is not a good predictor of the presence of a light anterior coloration in the other populations, in which this character seems to be environmentally driven (Fig. 8c). The proposed intergradation between *V*. *t*. *tristis* and *V*. *t*. *orientalis* (Pianka 2004) is most likely the result of confusion regarding color variation in CT + SE + SW + TE + WS (see below). On the other hand, the paraphyly of *V*. *t*. *orientalis* recovered by Fitch et al. (2006) is the result of labelling an individual from Noonbah Station, Queensland, as *V*. *t*. *orientalis*. In our study, both mitochondrial and nuclear data recovered that individual as a member of SE. Thus, multiple lines of evidence consistently indicate that NE is largely or fully isolated reproductively.

The distribution of *V*. *t*. *orientalis* suggests that the Great Dividing Range played a major role in its divergence from *V*. *t*. *tristis*. However, the uplift of this mountain range started roughly 70 Ma (Taylor 1994), while we estimated *V*. *t*. *orientalis* to have diverged about 2.99 Ma during the Pliocene. Furthermore, portions of the Great Dividing Range consist of low-elevation rolling hills (Johnson 2009), which are unlikely to represent a barrier to gene flow. However, the Range creates a heterogeneous mosaic of climatic conditions and vegetation types (Taylor 1994). This heterogeneity is probably accentuated during periods of environmental change, fragmenting suitable habitats. A period of increased aridity and cooling is inferred to have occurred during the Pliocene (Bowler 1982). This had an impact on genetic structure in other taxa (Garrick et al. 2004) and provides a likely explanation for the divergence of *V*. *t*. *orientalis*.

Our results show that *V*. *t*. *tristis* comprises several populations connected by gene flow (Figs. 3a, 5c). Pleistocene climatic fluctuations appear to be primarily responsible for genetic structure (Fig. 5b). The GDM suggests that climate accounts for more genetic variation during cooler periods such as the LGM (Fig. 4c). These time periods in Quaternary Australia are characterized not just by lower temperatures, but also by increased aridity, expansion of dune fields, and counterintuitively by the widespread presence of water bodies as a result of increased runoff (Pepper and Keogh 2021). All of these factors probably contributed to the fragmentation of the range of *V*. *t*. *tristis*. SE appears to be the most highly differentiated population. The Carpentarian Gap, Simpson Desert, and Lake Eyre are all major barriers that probably restrict gene flow between SE and other populations (Austin et al. 2013; Pepper and Keogh 2021; Pavón-Vázquez et al. 2022). However, occasional gene flow is apparently possible (Fig. 3, 5c). We also found that mesic areas that may have functioned as refugia in the face of increasing aridity harbor endemic lineages (Pepper et al. 2011; Catullo et al. 2014; Dalmaris et al. 2015; Pepper and Keogh 2021). These are the Top End region in the north, the Mediterranean regions in the southwest, and the Musgrave Ranges in central Australia (Figs. 1b, 2a). While these populations appear to be isolated from each other, they all show signs of admixture with a lineage that is widespread in the Arid Zone (Fig. 2a). A similar pattern was also found in another Australian monitor lizard (Pavón-Vázquez et al. 2022). This suggests that the expansion of arid-adapted lineages may have prevented or reversed speciation across several Australian taxa.

The variation in color exhibited by *V*. *t*. *tristis* is noteworthy and likely contributed to the taxonomic misperception in the group. To further add to the confusion, the type localities of *Monitor tristis* Schlegel 1839 and *Varanus punctatus* var. *orientalis* Fry 1913 are on opposite ends of the continent (some 3,500 km away) and comparative material was scarce until the second half of the 20^th^ century. In this study, we found that individuals assignable to SW are almost completely black (Schmida 2020), light-headed individuals are more prevalent in the Monsoonal Tropics but are also prevalent in the Musgrave Ranges, and widespread populations like WS and SE host both light and dark-headed individuals (Fig. 8b). Our analyses indicate that color variation in *V*. *t*. *tristis* is probably not linked to gene flow with *V*. *t*. *orientalis* and is instead driven by environmental conditions. Particularly, it follows a well-established pattern where darker colorations are expected to be more common farther from the Equator (Gaston et al. 2008). This is termed the thermal melanism hypothesis, as it is thought that darker colors are more prevalent in colder climates because they allow individuals to gain heat faster (Clusella Trullas et al. 2007). This pattern is expected to be more prevalent in ectotherm organisms that rely largely on external sources of heat (Bogert 1949). While we did not include temperature in our main analyses because it is tightly correlated with latitude, we confirmed that the probability of being dark-headed is negatively correlated with annual mean temperature (Fig. S7). In fact, the latter variable accounts for more variation in color than any other tested predictors except for latitude (Table S11).

Intriguingly, we found that the probability of being dark-headed is higher in areas with higher annual mean solar radiation (Fig. 8c). However, this predictor fails to explain much of the color variation (Table S11) and the 95% credible intervals of the conditional effects are wide (Fig. 8c).

### Implications for other Varanus

The nuclear and mitochondrial data indicate that *V*. *scalaris* WT is more closely related to *V*. *glauerti* and *V*. *tristis* than to other *V*. *scalaris*. Additionally, the first author was able to examine 20 specimens assignable to *V*. *scalaris* WT. The taxon shows a cluster of enlarged scales on each side of the vent (spinose in adult males), a trait shared with *V*. *glauerti* and *V*. *tristis* and not found in *V*. *scalaris*. On the other hand, *V*. *scalaris* WT can be distinguished from *V*. *glauerti* by having a relatively shorter tail in adults (< 65% of total length), and from *V*. *tristis* by having the distal end of the tail banded. Thus, *V*. *scalaris* WT likely represents an independent species within the *Odatria* subgenus. We provide a distribution map for this lineage in Fig. S8.

Additional sampling is necessary to confidently evaluate the taxonomic status of *V*. *glauerti* from the Top End. Morphologically, this population is clearly assignable to *V*. *glauerti* and distinct from *V*. *tristis*. Still, our mitochondrial tree suggests that this population is sister to *V*. *t*. *tristis*. Brennan et al. (2021) recovered similar results based on hundreds of nuclear loci. However, they only included the same single sample present in our mitochondrial tree. In the absence of additional data, it is difficult to evaluate alternative explanations for the phylogenetic placement of our single sample of *V*. *glauerti* from the Top End such as introgression or incomplete lineage sorting.

## Conclusions

Our study shows how genomic data and the information provided by museum specimens and community observations can be integrated to shed light on species limits. We analyzed these data through a hypothesis-driven workflow in a variable species complex. With human activities rapidly accelerating extinction (Barnosky et al. 2011), it is urgent to promptly but accurately characterize biodiversity. Community science and novel methods for the fast processing and analysis of data (such as machine learning) are likely to streamline species delimitation, allowing faster progress than that possible under traditional taxonomic practices. These strategies are likely to have a greater impact on taxonomic research in the Global South, where most megadiverse countries are located (Prathapan and Dharma Rajan 2020). Embracing these approaches may lead us into a more productive and equitable new era of biodiversity research based on global cooperation between communities and scientists.

## Acknowledgments

We thank A.P. Amey, R.D. Bray, P.D. Campbell, P.J. Couper, G.M. Dally, A. Drew, M.R. Hutchinson, L. Joseph, C. Kovach, S. Mahony, J. Melville, J.J.L. Rowley, J.W. Streicher, S. South, J. Sumner, A. Velasco Castrillón, and J. Worthington Wilmer for the support provided during specimen examination in their respective institutions and for the loan of tissue samples; M. Arvizu Meza for her help with specimen examination; R.J. Ellis, J.E. Farquhar, K. Palmer, W. Scott, and S.M. Zozaya for sharing relevant photographs; D. Esquerré and J. Fenker for methodological suggestions; and A. Kilian and M. Pepper for their help with sequencing.

