## Supplementary Figures for "Integrating Genomics, Collections, and Community Science to Reveal Speciation in a Variable Monitor Lizard (*Varanus tristis*)"

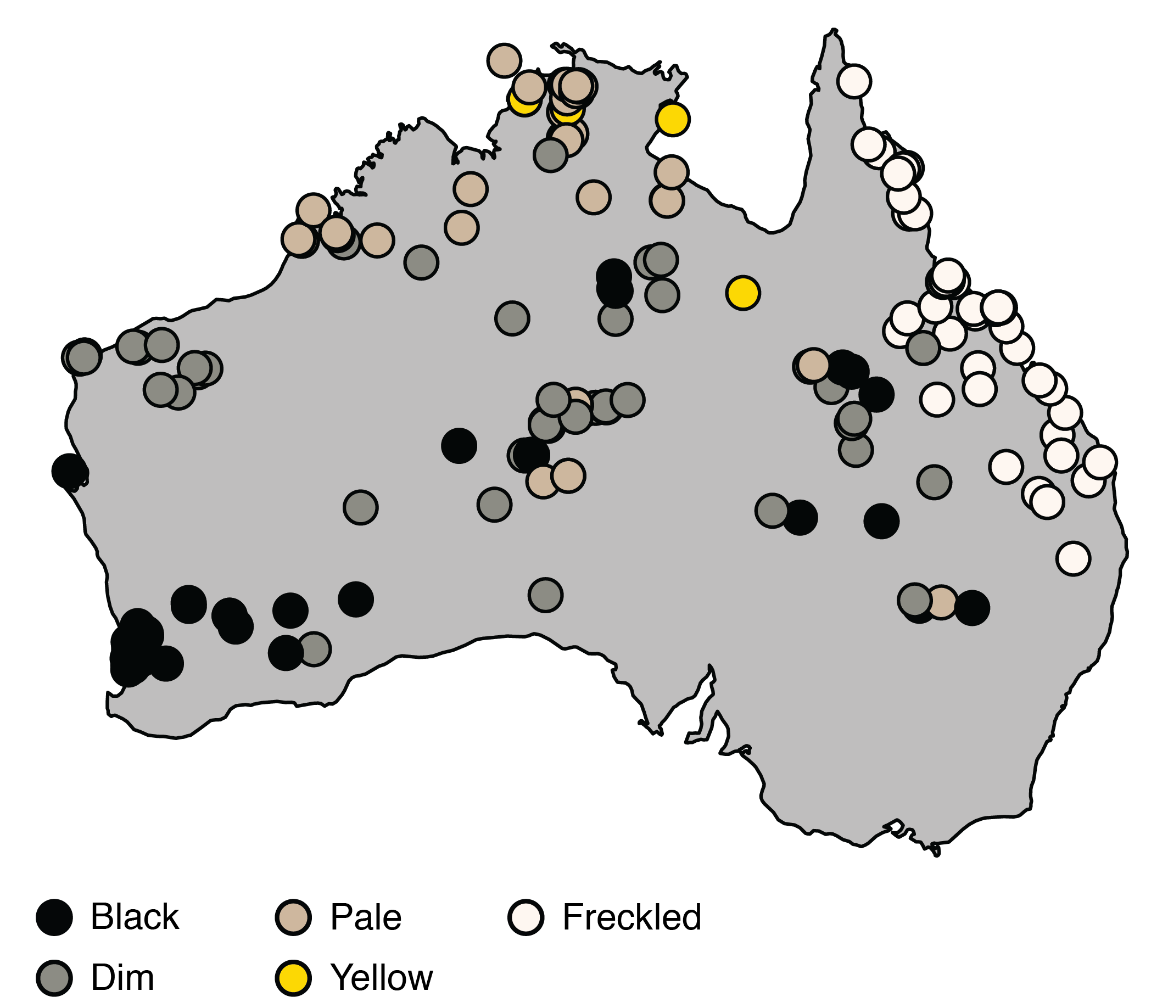

Figure S1. Distribution of color variation classified into five discrete categories (see Fig. 8).

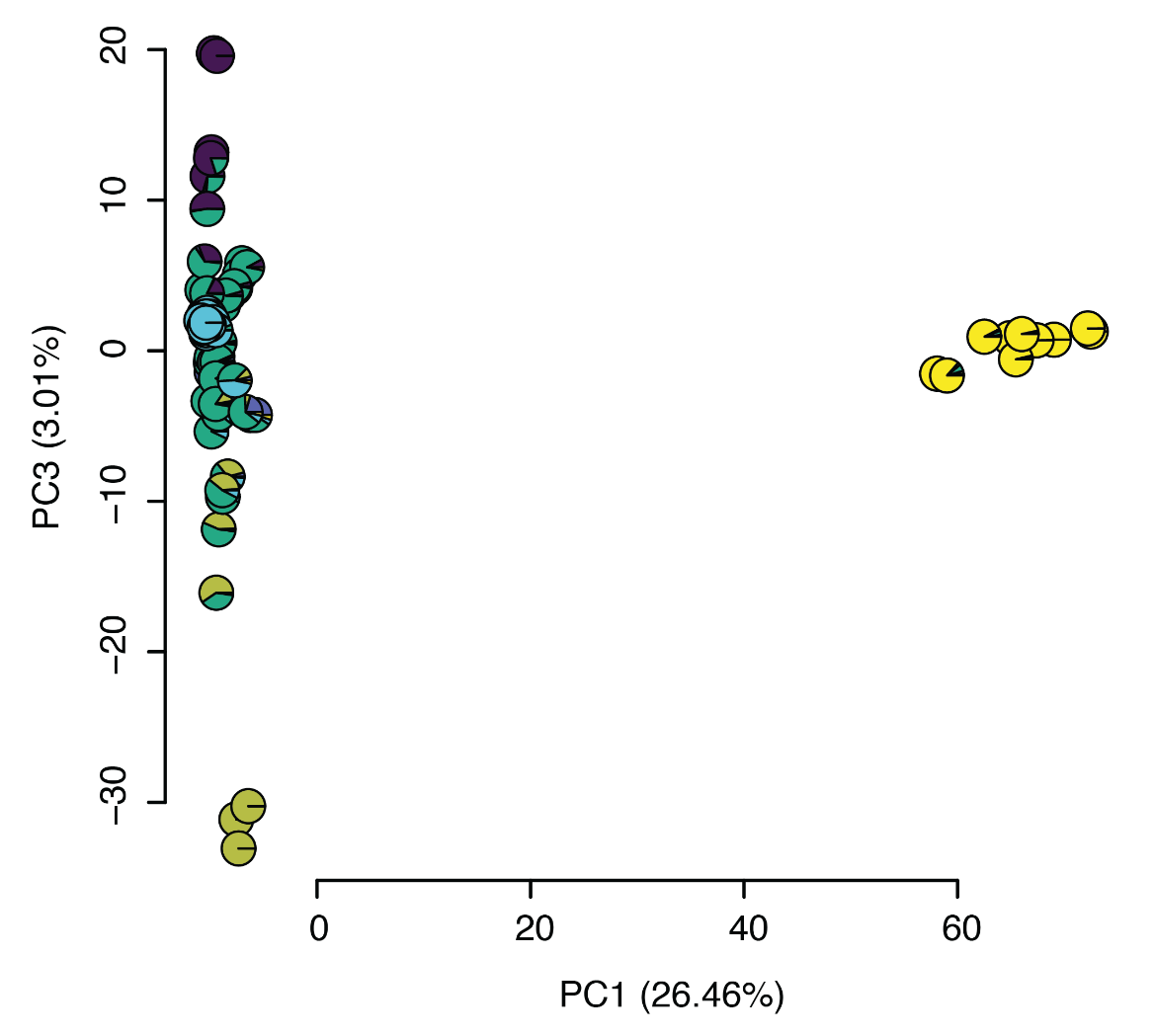

Figure S2. Principal components one and three based on the analysis of the SNP data. Pie charts indicate ancestry coefficients for individuals. Colors follow Fig. 3.

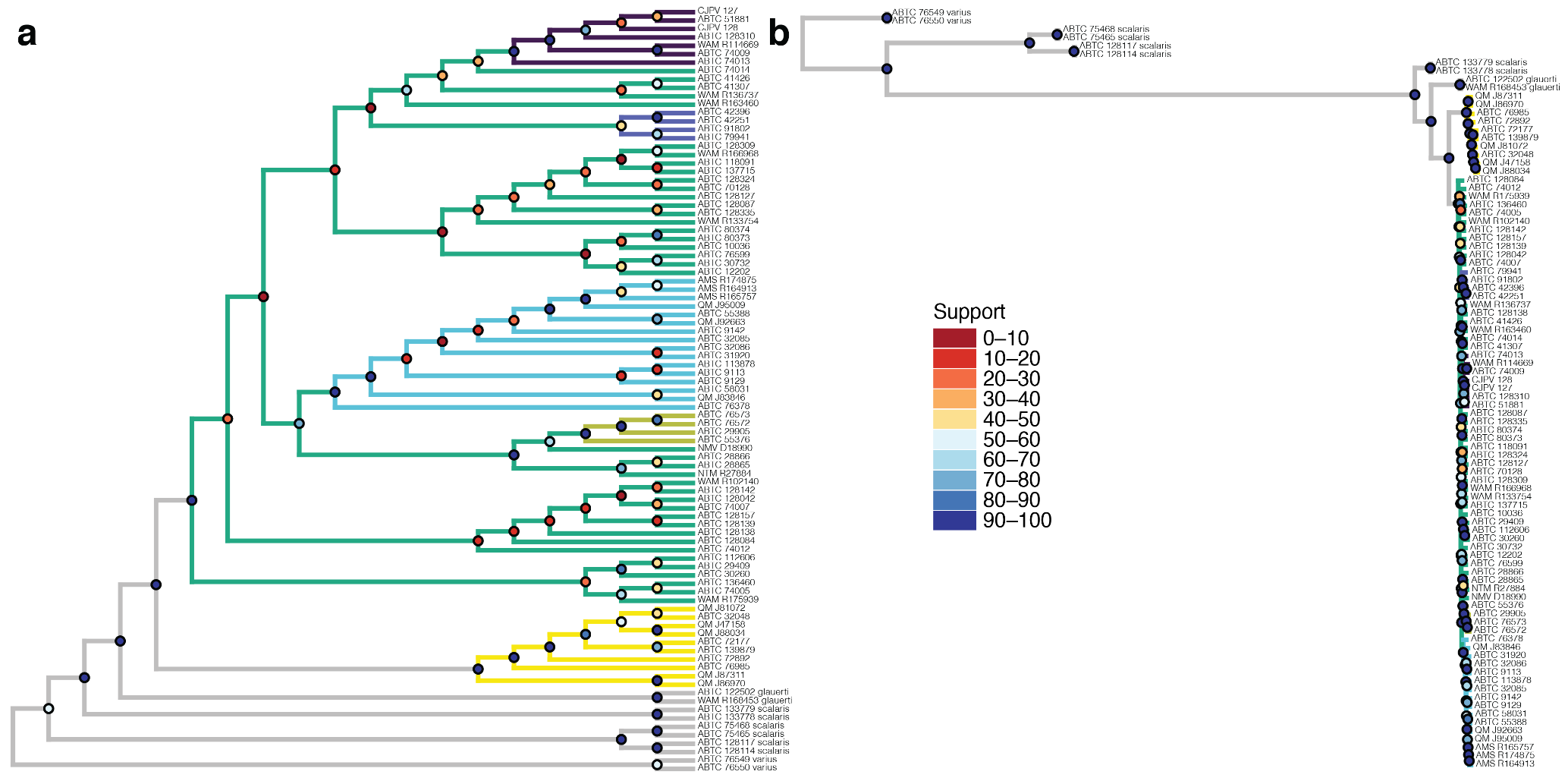

Figure S3. Nuclear phylogeny of *V*. *tristis*. Branch colors follow Fig. 5a. a) Individual-level phylogeny estimated with SVDquartets; circles at nodes indicate bootstrap support; branch lengths are arbitrary. b) Individual-level phylogeny estimated with IQ-TREE; circles at nodes indicate ultrafast bootstrap support; branch lengths are proportional to the number of substitutions per site.

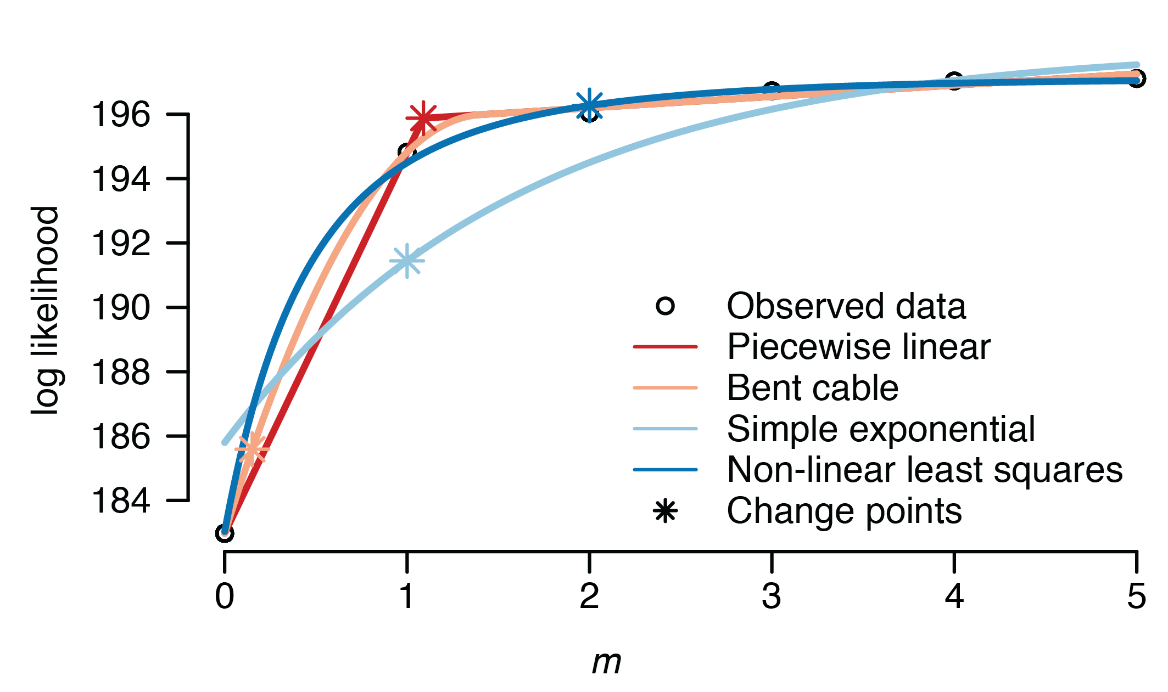

Figure S4. Support for different number of migration edges (*m*) in the TreeMix analyses under different models.

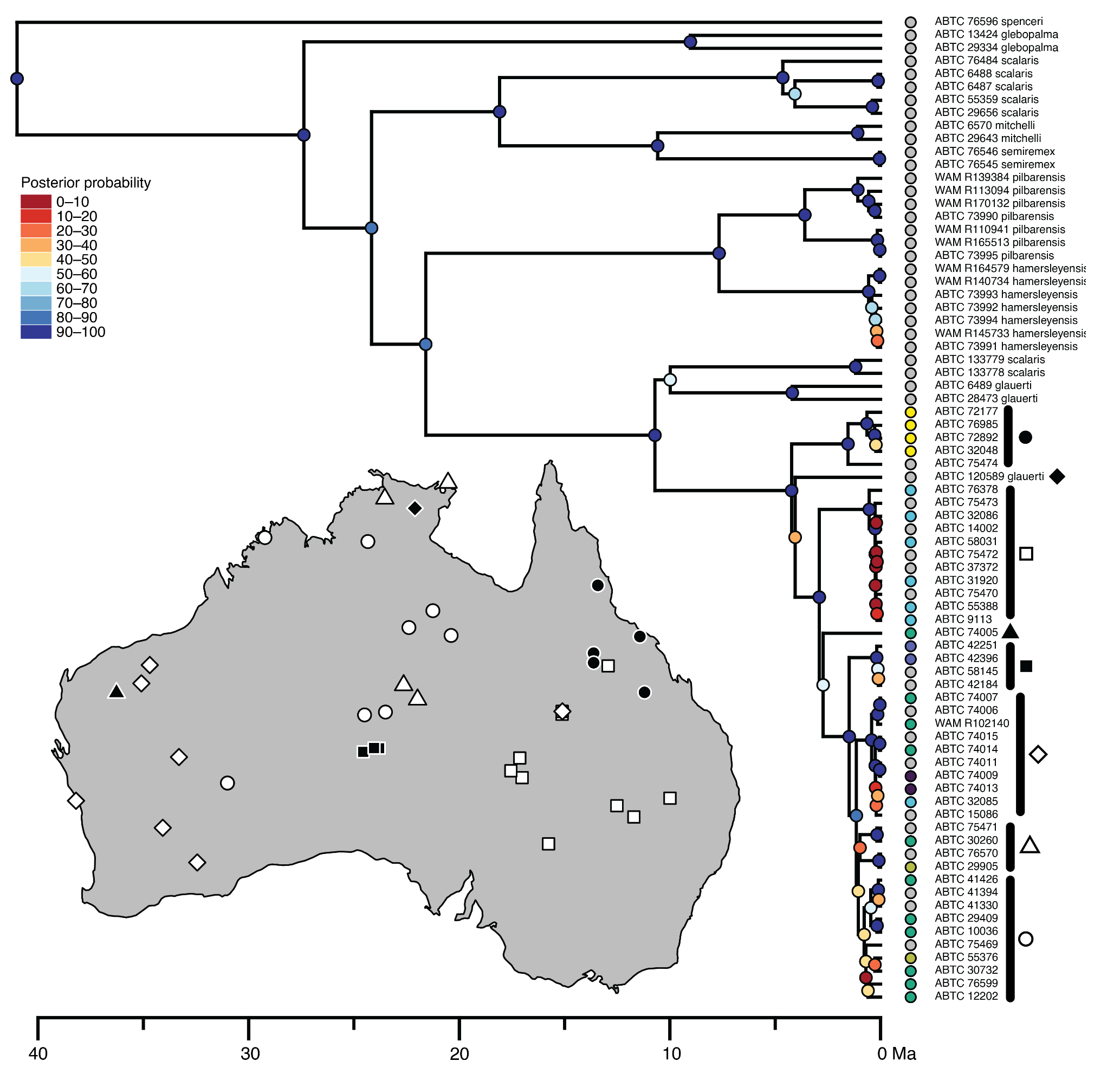

Figure S5. Time-calibrated Bayesian mitochondrial phylogeny obtained with BEAST. Circles next to tips indicate the population with the highest nuclear ancestry coefficient for each tip estimated with sNMF (colors follow Fig. 3). The map indicates the distribution of major mitochondrial clades of *V*. *tristis*.

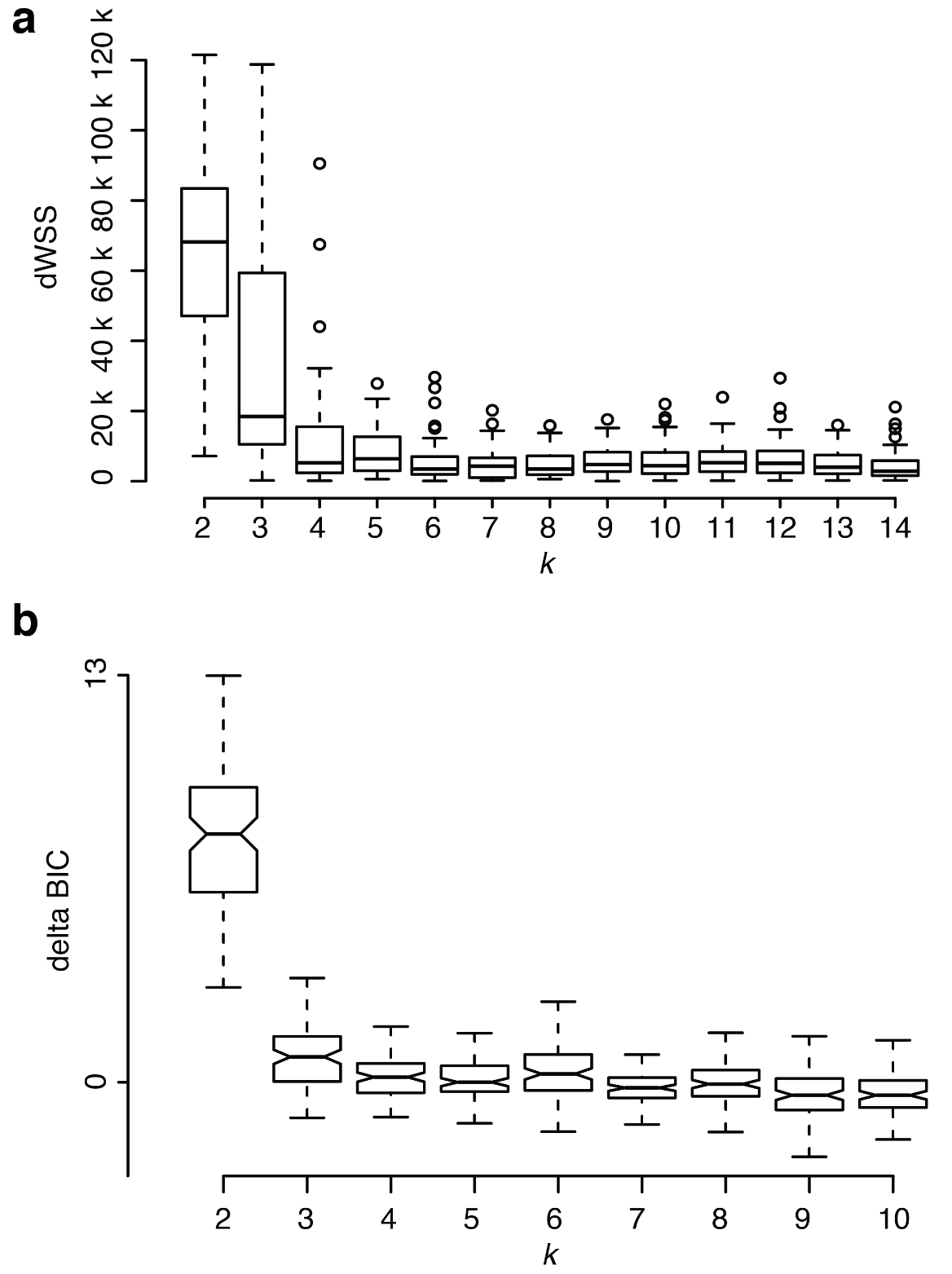

Figure S6. Support for different number of clusters (*k*) in the SOM and SuperSOM analyses. a) Support based on the decrease in the weighted sum of squares (dWSS) across 50 replicates for the analyses based on the SNP data. b) Support based on the Bayesian Information Criterion (BIC) across 100 replicates for the analyses based on the combined data.

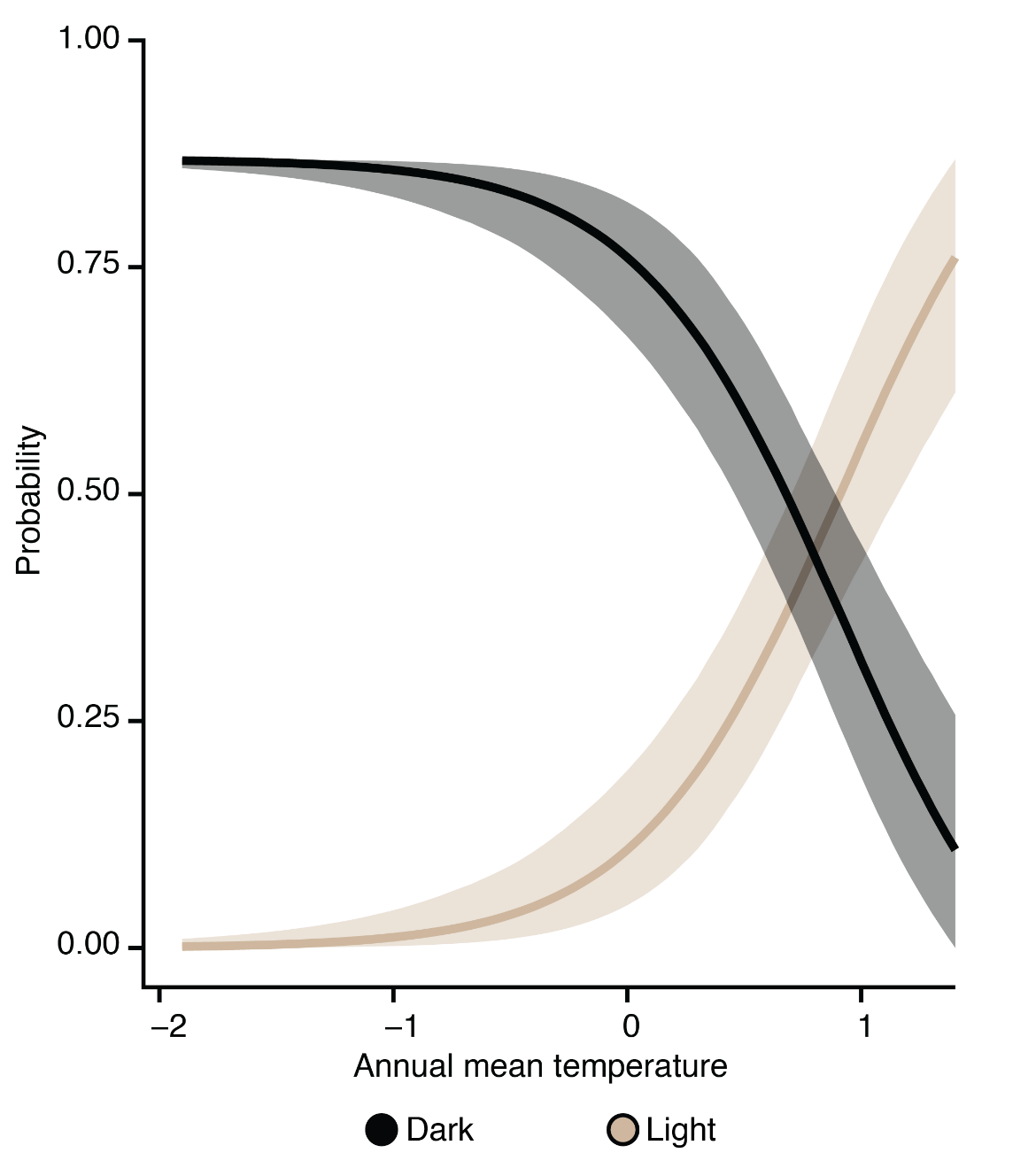

Figure S7. Conditional effects describing the relationship between color and annual mean temperature in *V*. *t*. *tristis*. The probability of being light or not was modelled; the probability of being dark was extrapolated from the former. Shaded contours indicate 95% credible intervals. Annual mean temperature was log-transformed and scaled.

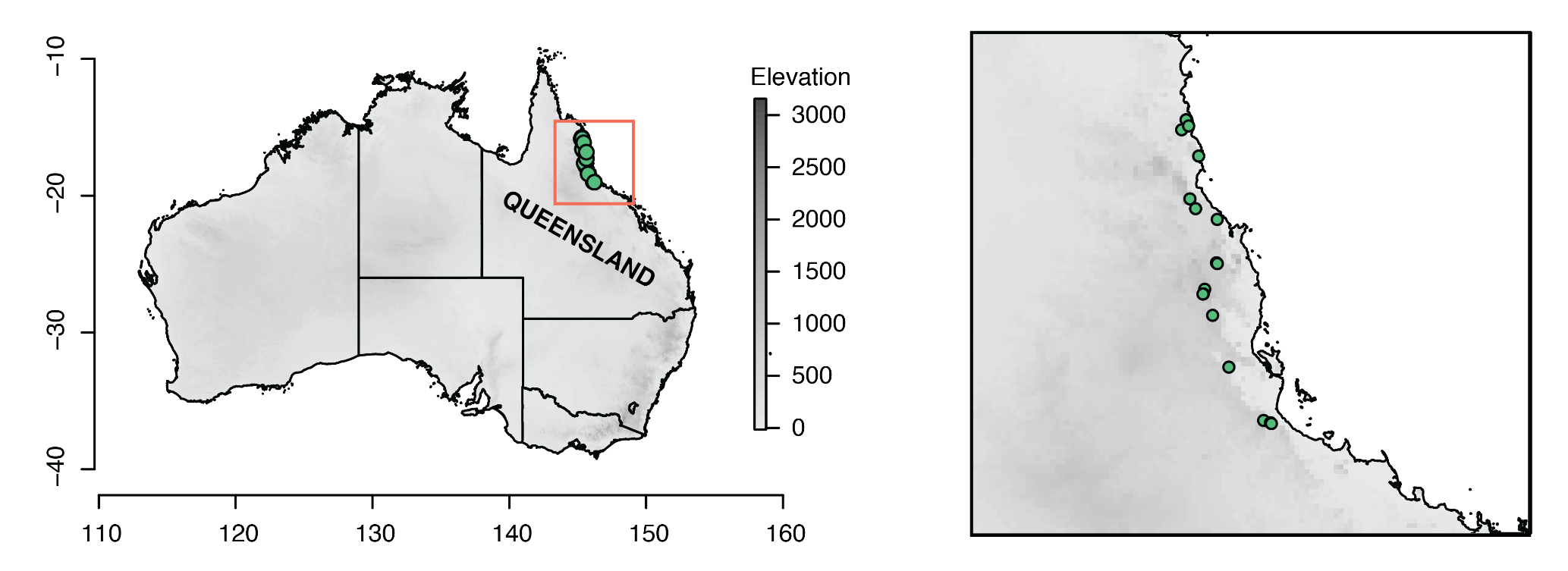

Figure S8. Distribution of examined specimens assigned to *V*. *scalaris* WT. The right panel shows a magnified version of the area inside the red square in the left panel.
